# Luminal nutrients activate distinct patterns in submucosal and myenteric neurons in the mouse small intestine

**DOI:** 10.1101/2021.01.19.427232

**Authors:** C. Fung, M.M. Hao, Y. Obata, J. Tack, V. Pachnis, W. Boesmans, P. Vanden Berghe

**Author notes:** Correspondence: Pieter Vanden Berghe.

## Abstract

Nutrient signals sensed by enteroendocrine cells are conveyed to the enteric nervous system (ENS) to initiate intestinal reflexes. We addressed whether there are specific enteric pathways dedicated to detecting different luminal nutrients. Calcium imaging was performed on intact jejunal preparations from Wnt1-cre;R26R-GCaMP3 and Villin-cre;R26R-GCaMP3 mice which express a fluorescent calcium indicator in their ENS or intestinal epithelium, respectively. Glucose, acetate, and L-phenylalanine were perfused onto the mucosa whilst imaging underlying enteric neurons. Nutrient transport or diffusion across the mucosa was mimicked by applying nutrients onto sensory nerve endings in a villus, or onto myenteric ganglia. The nutrients perfused onto the mucosa each elicited Ca^2+^ transients in submucosal neurons and in distinct patterns of myenteric neurons. Notably, the neurochemical subtypes of myenteric neurons that responded differed between the nutrients, while submucosal responders were predominantly cholinergic. Nutrients applied into villi or onto ganglia did not elicit specific neuronal responses but did stimulate Ca^2+^ signaling in the mucosal epithelium. These data suggest that nutrients are first detected at the level of the epithelium and that the ENS is capable of discriminating between different compositions of luminal content. Furthermore, our data show that responses to mucosal stimulation are primarily in the myenteric plexus and submucosal neurons respond secondarily.

## Introduction

Monitoring of ingested nutrients by the gastrointestinal (GI) tract is essential for maintaining the body’s energy homeostasis. The gut contains its own neural network, that is the Enteric Nervous System (ENS), which comprises a complete repertoire of intrinsic primary afferent neurons (IPANs, intrinsic sensory neurons with Dogiel II morphology), interneurons, and motor neurons organized within two ganglionated plexuses: the submucosal and myenteric plexus (Furness, 2012, Schneider et al., 2019, Fung and Vanden Berghe, 2020, Li et al., 2020, Spencer and Hu, 2020). The ENS is situated solely within the walls of the GI tract. Thus, it is well positioned and equipped to sample the local intestinal environment and respond accordingly by modulating intestinal secretion, blood flow, and motility. The presence of luminal nutrients can trigger local enteric nerve reflexes (Gwynne and Bornstein, 2007a) and subsequently evoke changes in intestinal motility (Gwynne et al., 2004, Gwynne and Bornstein, 2007b, Ellis et al., 2013). In addition to nutrients, there is a heightened interest in the role of gut microbiota residing in the lumen, particularly given its implications in a number of diseases ranging from obesity to neurodegenerative disease (Obata and Pachnis, 2016, Chalazonitis and Rao, 2018, Cryan et al., 2019). Despite this, how different luminal signals, even basic nutrients, are communicated to the ENS to trigger appropriate physiological responses within the intestine and the specific enteric pathways that are involved remain elusive.

Enteric nerves do not directly contact the luminal contents; rather, they are shielded from the external luminal environment by a single cell layer of mucosal epithelium. The mucosa is the initial site at which luminal contents are sensed and plays a dual role in the absorption of nutritious luminal contents and in acting as a protective barrier against harmful pathogens. Specialized sensor cells, namely enteroendocrine cells (EECs), sparsely scattered throughout the epithelium are important in monitoring the intestinal milieu and serve as an interface between the lumen and the ENS (Gribble and Reimann, 2016). There is an extensive body of literature on characterizing the molecular aspects of chemosensation at the level of the mucosa (Depoortere, 2014, Liddle, 2018), and it is clear that EECs contain a diverse range of signaling molecules such as 5-HT, GLP-1, and CCK (Diwakarla et al., 2017, Fothergill and

Furness, 2018). However, the specific mucosal mediators that signal to the ENS remain elusive. Thus, we lack an understanding of the fundamental link between the sensors that directly detect luminal contents, and the ENS, which triggers the gut to take any necessary actions.

Some EECs make specialized contacts with extrinsic sensory afferents via their ‘neuropod’ (Kaelberer et al., 2018), providing a direct link between the luminal environment and the central nervous system and a means of rapidly detecting nutrient information. Whether similar connections between EECs and enteric neurons exist and whether there are specific enteric pathways for detecting different luminal nutrients remain unclear. Due to the complex spatial organization of the innervated epithelium, lamina propria and underlying nerve plexuses, studying their functional connectivity while maintaining the 3D architecture of the intact gut wall has traditionally been technically challenging. Indeed, our current knowledge of enteric responses to chemical or mechanical stimulation of the mucosa has been gleaned from various studies which used preparations where the mucosa was partially dissected in order to access and study the electrophysiological responses in the exposed neurons (Bertrand et al., 1997, Bertrand et al., 1998, Bertrand et al., 2000, Gwynne and Bornstein, 2007a). To further our understanding of how luminal contents are perceived by the ENS, it is necessary to preserve the integrity of all the gut layers and the connections between them. This can now be tackled optically with the availability of transgenic mice that selectively express fluorescent activity-dependent reporters in their ENS (Boesmans et al., 2017, Hao et al., 2020).

In this study, we examined how luminal information is communicated from the mucosa to the ENS using full thickness preparations from the jejunum of Wnt1xGCaMP3 and VillinxGCaMP3 mice. We aimed to first address whether there are specific enteric circuits dedicated to detecting different luminal contents such that the ENS can discriminate between different food compositions and act accordingly. By analogy with the olfactory system, where most odors are not encoded by a single receptor but by a pattern of receptors and olfactory receptor neurons (Su et al., 2009), different luminal stimuli may similarly be identified by the activation of different patterns of enteric neurons. Our results indicate that different luminal nutrients can elicit patterned Ca^2+^ responses in specific myenteric neurons and that luminal information is first transmitted to the myenteric plexus and then to the submucosal plexus.

## Results

### Mucosal depolarization activates a specific subset of enteric neurons

Under physiological conditions, the gut epithelium is exposed to a plethora of nutrient and chemical stimuli that widely activates many EECs including 5-HT-containing enterochromaffin (EC) cells (Martin et al., 2017b, Martin et al., 2017a, Gribble and Reimann, 2019). High K^+^ (75 mM) solution was used as a means of broadly stimulating these EECs which are electrically excitable (Rogers et al., 2011, Bellono et al., 2017) and to examine the neurons that are subsequently activated in the underlying submucosal or myenteric plexus of intact intestinal preparations from Wnt1 |GCaMP3 mouse jejunum (Figure 1). Mucosal depolarization evoked Ca^2+^ transients in 24 ± 6% of submucosal neurons (n = 132 neurons, N = 7 preparations) and 22 ± 3% of myenteric neurons (n = 175, N = 5). To further characterize the responding submucosal neurons, the imaged preparations were labelled *post hoc* for choline acetyltransferse (ChAT) and vasoactive intestinal peptide (VIP). These markers distinguish between the 2 major and distinct subtypes of secretomotor/vasodilator neurons in the submucosal plexus (Mongardi Fantaguzzi et al., 2009). In the submucosal plexus, responding neurons were nearly all cholinergic (ChAT^+^; 13/14 neurons, N = 4; Figure 1B-D”; Table 2), and rarely VIP^+^ (1/14 neurons). For myenteric plexus preparations, we examined calbindin- and nNOS-immunoreactivity which are commonly used markers for intrinsic primary afferent neurons (IPANs) and inhibitory inter- and motor neurons, respectively (Qu et al., 2008). The majority of myenteric neurons that responded were calbindin^+^ (18/22 neurons, N = 4; Figure 1F-H”; Table 3) and only few were nNOS^+^ (2/22 neurons), consistent with the notion that predominantly IPANs were activated.

**Figure 1:**
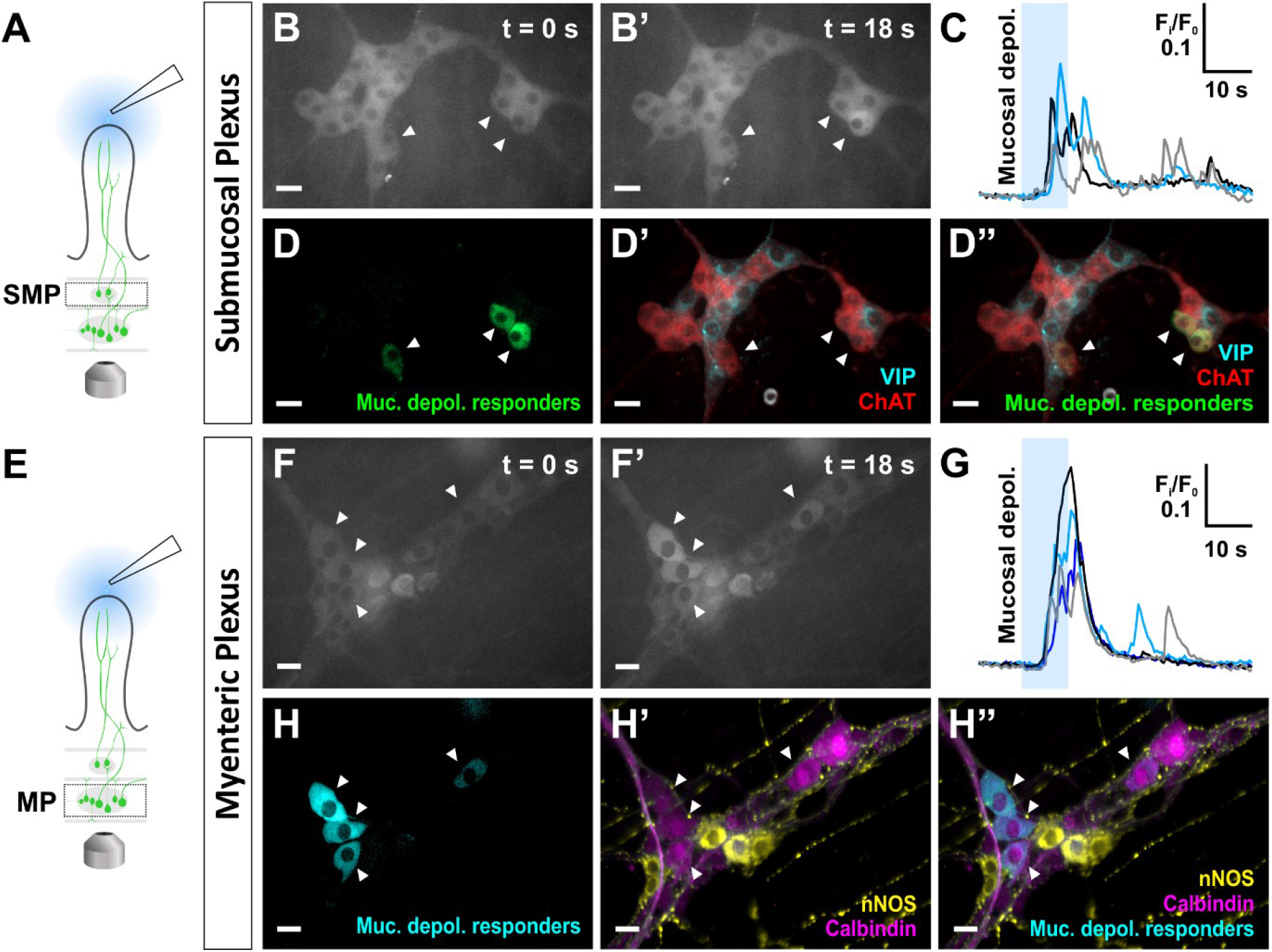
Enteric neuronal responses to mucosal depolarization in full thickness preparations of Wnt1|GCaMP3 mouse jejunum. **A.** The mucosa was depolarized by perfusing a high K^+^ (75 mM) Krebs solution onto the surface from t = 10 s to t = 20 s while imaging the underlying submucosal plexus. **B-B’.** A subset of submucosal neurons respond to mucosal depolarization as marked by the white arrowheads. **C.** Calcium transients of the responding submucosal neurons indicated in **B-B’**. **D.** Mucosal depolarization responders are shown in green. **D’.** The corresponding ganglion immunostained for the cholinergic marker choline acetyltransferase (ChAT) and vasoactive intestinal peptide (VIP). **D”.** An overlay of panels **D** and **D’** demonstrate that responders to mucosal depolarization were cholinergic (ChAT^+^) submucosal neurons. **E.** The mucosa was depolarized by perfusing a high potassium (75 mM) Krebs solution onto the surface from t = 10 s to t = 20 s while imaging the underlying myenteric plexus. **F-F’.** A subset of myenteric neurons respond to mucosal depolarization (marked by the white arrowheads). **G.** Calcium transients of the responding myenteric neurons indicated in **F-F’**. **H.** Mucosal depolarization responders are depicted in cyan. **H’.** The corresponding ganglion immunolabeled for calbindin and neuronal nitric oxide synthase (nNOS). **H”.** An overlay of panels **H** and **H’** demonstrate that responders to mucosal depolarization were calbindin^+^ myenteric neurons. Scale bars = 20 μm.

We then assessed whether intact nerve projections from the myenteric plexus to the mucosa are necessary for the transmission of the luminal signal evoked by mucosal depolarization by severing these connections. First, we established that mucosal depolarization elicits Ca^2+^ responses in myenteric neurons in intact preparations (n = 93, N = 3; Figure 2A-C). The gut layers were then peeled apart to severe the connections between the myenteric plexus and mucosa, and the separate layers were placed back together. This procedure completely abolished the myenteric responses to mucosal depolarization in ganglia previously imaged (Figure 2D-F), demonstrating that myenteric projections to the mucosa are indeed essential for communicating the luminal signal. In addition, this shows that the mucosal barrier remains intact in our preparations. Furthermore, to confirm that the neurons were still viable following peeling, we then showed that directly depolarizing the same ganglia evoked robust Ca^2+^ transients in nearly all neurons and includes all of those that initially responded (89/93 neurons; Figure 2G-I).

**Figure 2:**
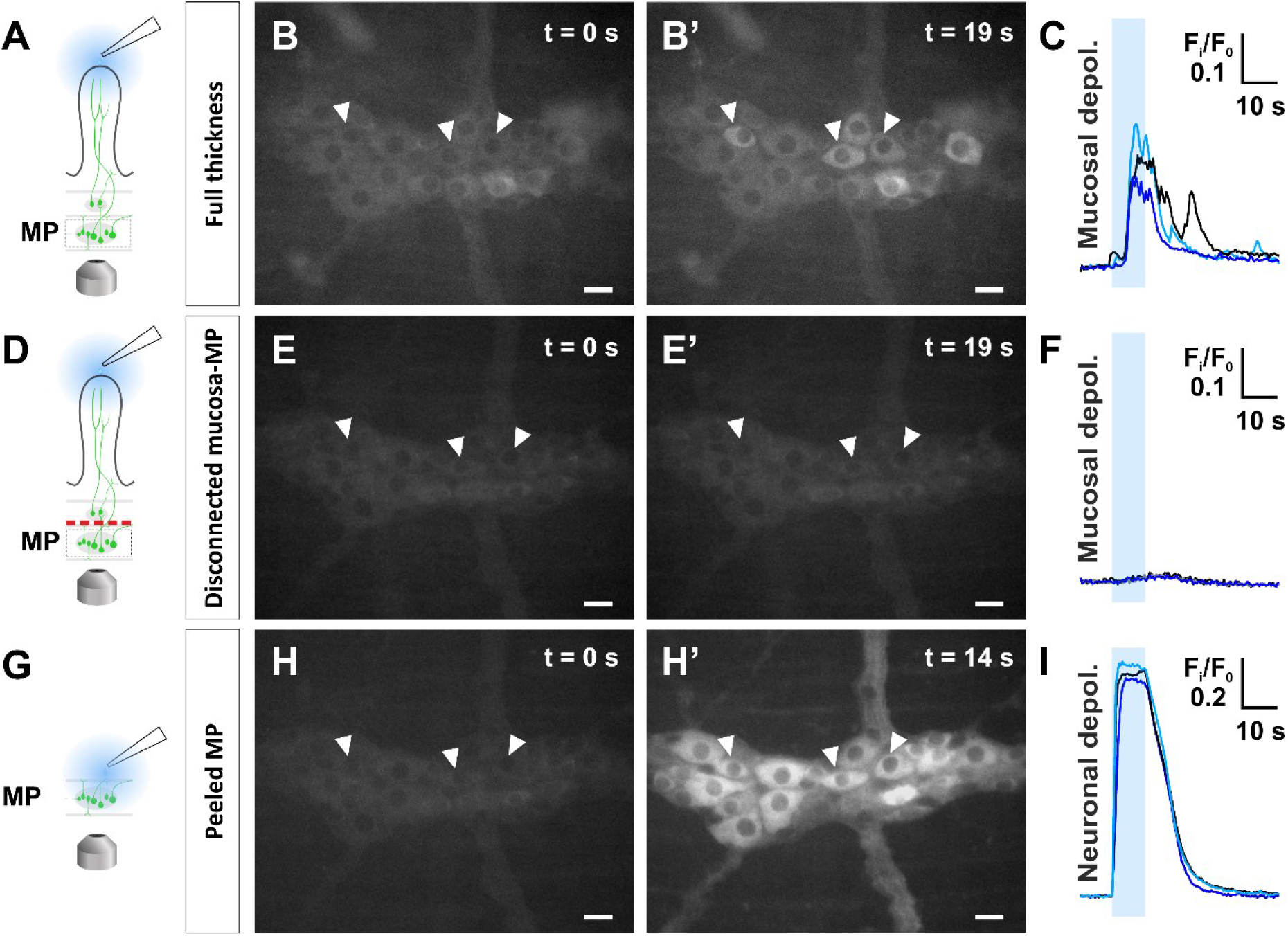
Intact nerve connections between the myenteric plexus and mucosa are necessary to transmit the response to mucosal depolarization. **A.** Myenteric responses to mucosal depolarization with high K^+^ (75mM) Krebs were first examined in full thickness preparations. **B-B’.** Select responding neurons are indicated with white arrowheads. **C.** Corresponding calcium transients of the responding neurons marked in the previous panels. High K^+^ Krebs was applied from t = 10 s to t = 20 s. **D.** The previously imaged preparations were peeled apart between the circular muscle and submucosal plexus layers to severe the connections between the myenteric plexus and mucosa. The separate layers were replaced and the mucosa was depolarized again. **E-F.** The same ganglion was imaged and responses to mucosal depolarization were abolished following peeling of the preparation. **G.** The mucosal layer was removed to expose the myenteric plexus and the neurons were directly exposed to the depolarizing K^+^ solution. **H-I.** All myenteric neurons in the previously imaged ganglion, including the initial responders, were depolarized. Scale bars = 20 μm.

### Different nutrients applied to the mucosa activate submucosal and myenteric neurons in distinct patterns

To determine whether different luminal nutrients can stimulate distinct patterns of enteric neurons, nutrient solutions were applied to the mucosa while imaging the activity in either the underlying submucosal plexus (Figure 3) or myenteric plexus (Figure 4). Glucose (300 mM; Suppl. Movies 1 and 2), acetate (100 mM; Suppl. Movies 3 and 4), and L-phe (100 mM; Suppl. Movies 5 and 6), were selected as a model sugar, short chain fatty acid, and amino acid, respectively.

**Figure 3:**
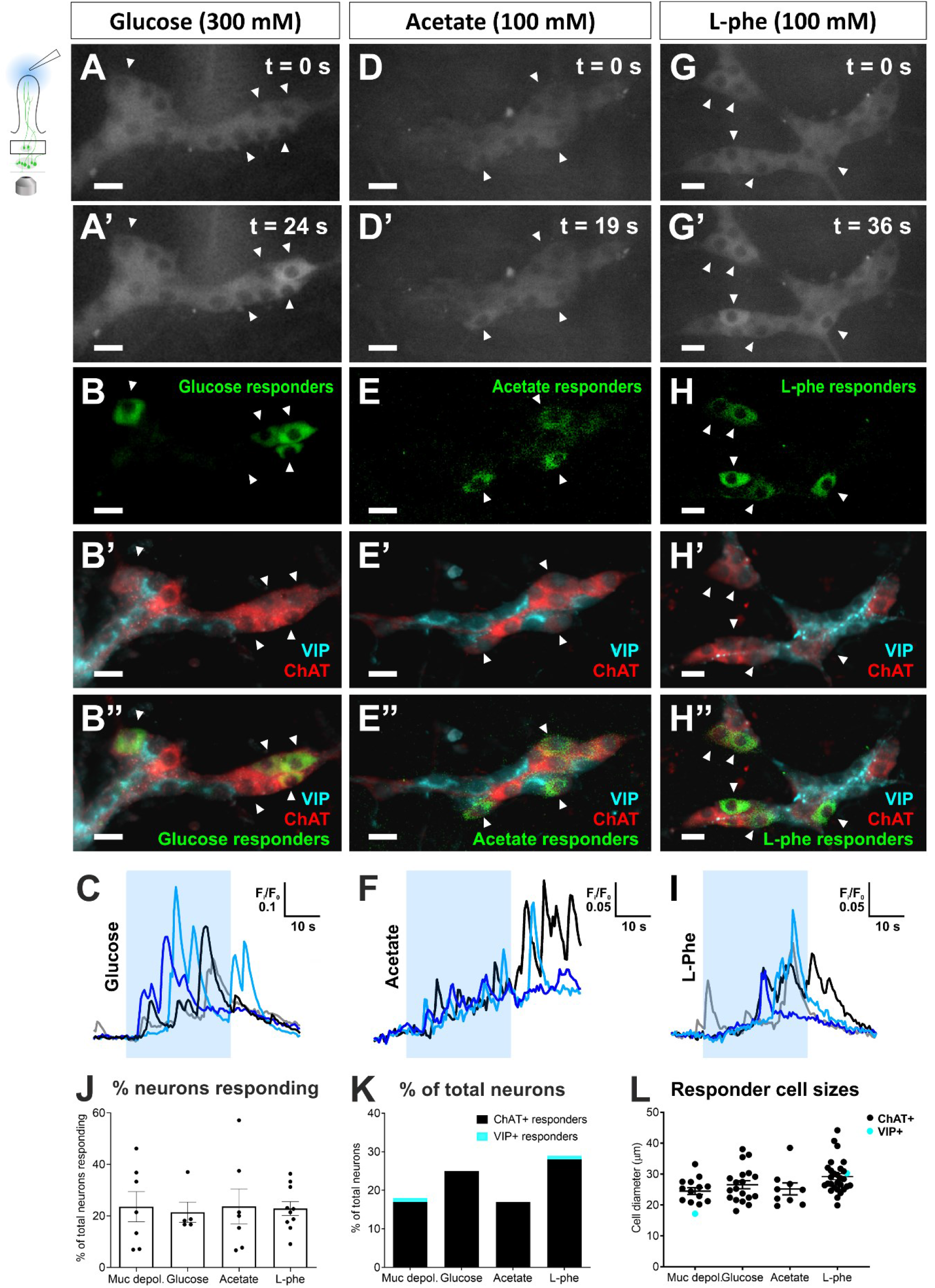
Submucosal responses to nutrients perfused onto the mucosa. **A-A’.** Mucosal perfusion of glucose (300 mM) activates a subset of submucosal neurons as marked by white arrowheads. Glucose was applied from t = 10 s to t = 40 s. Scale bars = 20 μm. **B.** Glucose responders are represented in green. **B’.** The corresponding ganglion labelled for choline acetyltransferase (ChAT) and vasoactive intestinal peptide (VIP). **B”.** An overlay of panels **B** and **B’** demonstrate that glucose responders were cholinergic (ChAT^+^) submucosal neurons. **C.** Corresponding calcium transients in the responding neurons marked in the previous panels. **D-D’.** Mucosal perfusion of acetate (100 mM) activates a subset of submucosal neurons as marked by white arrowheads. Acetate was applied from t = 10 s to t = 40 s. **E.** Acetate responders are shown in green. **E’.** The corresponding ganglion immunolabelled for ChAT and VIP. **E”.** An overlay of panels **E** and **E’** demonstrate that acetate responders were cholinergic (ChAT^+^) submucosal neurons. **F.** Corresponding calcium transients in the responding neurons marked in the previous panels. **G-G’.** Mucosal perfusion of L-phe (100 mM) activates a subset of submucosal neurons as marked by white arrowheads. L-phe was applied from t = 10 s to t = 40 s. **H.** L-phe responders are depicted in green. **H’.** The corresponding ganglion stained for ChAT and VIP. **H”.** An overlay of panels **H** and **H’** demonstrate that L-phe responders were ChAT^+^ submucosal neurons. **I.** Corresponding calcium transients in the responding neurons marked in the previous panels. **J.** The percentage of neurons within the field of view that responded to mucosal depolarization, glucose, acetate, and L-phe were comparable. **K.** The majority of responders to each stimulus tested were cholinergic (ChAT^+^). **L.** The cell diameter of the responders to each stimulus were generally similar, although a significant difference was detected between responders to mucosal depolarization and L-phe (One-way ANOVA, Tukey’s multiple comparisons test, *P* < 0.05). The cell diameter of responders was comparable between different nutrients.

**Figure 4:**
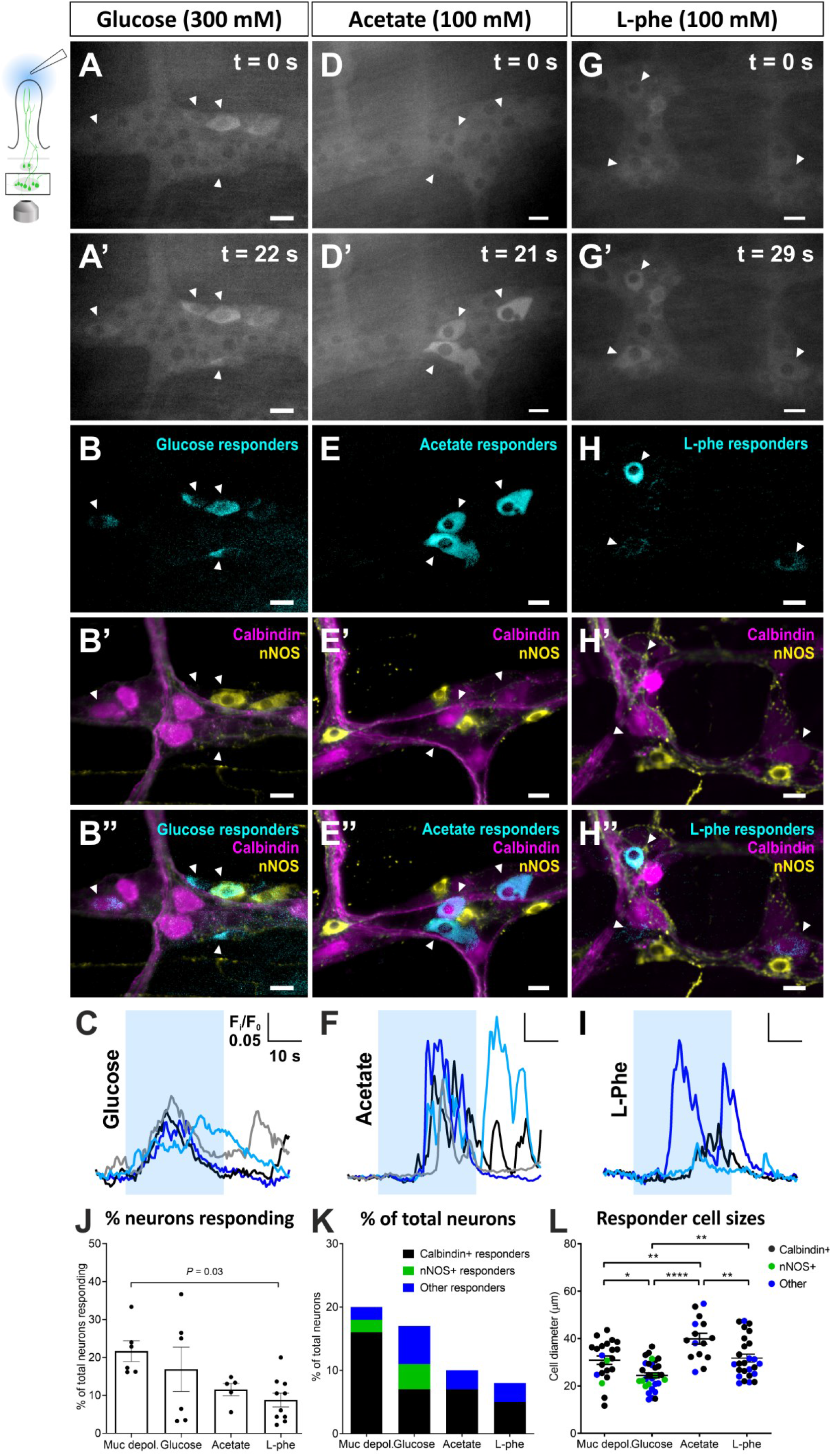
Myenteric responses to nutrients perfused onto the mucosa. **A-A’.** Mucosal perfusion of glucose (300 mM) activates a subset of myenteric neurons as marked by white arrowheads. Glucose was applied from t = 10 s to t = 40 s. Scale bars = 20 μm. **B.** Glucose responders are represented in cyan. **B’.** The corresponding ganglion following *post hoc* immunolabelling for the calbindin and neuronal nitric oxide synthase (nNOS). **B”.** An overlay of panels **B** and **B’** show that glucose responders include calbindin^+^ and nNOS^+^ myenteric neurons. **C.** Corresponding calcium transients in the responding neurons marked in the previous panels. **D-D’.** Mucosal perfusion of acetate (100 mM) activates a subset of myenteric neurons as marked by white arrowheads. Acetate was applied from t = 10 s to t = 40 s. **E.** Acetate responders are shown in cyan. **E’.** The corresponding ganglion labelled for calbindin and nNOS. **E”.** An overlay of panels **E** and **E’** demonstrate that many acetate responders were calbindin^+^. **F.** Corresponding calcium transients in the responding neurons marked in the previous panels. **G-G’.** Mucosal perfusion of L-phe (100 mM) activates a subset of myenteric neurons as marked by white arrowheads. L-phe was applied from t = 10 s to t = 40 s. **H.** L-phe responders are depicted in cyan. **H’.** The corresponding ganglion stained for calbindin^+^ and nNOS^+^. **H”.** An overlay of panels **H** and **H’** demonstrate that some L-phe responders were calbindin^+^ myenteric neurons. **I.** Corresponding calcium transients in the responding neurons marked in the previous panels. **J.** The percentage of neurons within the field of view that responded to mucosal depolarization, glucose, and acetate were comparable. Only L-phe evoked a response in a significantly smaller percentage of neurons compared to mucosal depolarization (One-way ANOVA, Tukey’s multiple comparisons test, *P* = 0.03). **K.** The majority of responders to each stimulus tested were calbindin^+^, while none of the acetate and L-phe responders were nNOS^+^. **L.** The cell diameter of the responders to each stimulus differed significantly (One-way ANOVA, Tukey’s multiple comparisons test, \**P* < 0.05; \*\**P* < 0.01; **** *P* < 0.0001).

In the submucosal plexus, each of the different nutrients applied evoked responses in a comparable proportion of neurons within the field of view. Glucose, acetate, and L-phe activated Ca^2+^ transients in 21 ± 4% (n = 106, N = 4), 19 ± 6% (n = 84, N = 6), and 19 ± 5% (n = 126, N = 9) of submucosal neurons, respectively (Figure 3). Notably, nearly all responding neurons were identified as cholinergic (ChAT^+^) neurons (Figure 3K; table 2). The size of cell bodies was used as another mode of classifying different enteric neurons, since IPANs tend to have type II morphology and cell bodies larger than other neurons (Qu et al., 2008). Thus, we also quantified the cell diameter of responding submucosal neurons and these were also comparable in cell size for each nutrient tested (Figure 3L).

In the myenteric plexus, glucose (300 mM) evoked Ca^2+^ transients in 17 ± 6% of myenteric neurons (n = 183, N = 6). The responding neurons were mainly calbindin^+^ (13/31) and some were neuronal nitric oxide synthase^+^ (nNOS^+^) (7/31 neurons), but many responders did not label for either marker (11/31 neurons) (N = 6; Figure 4A-C; table 3). Acetate (100 mM) elicited responses in 13 ± 2% myenteric neurons (n = 153, N = 6). Of the responding myenteric neurons, most were calbindin^+^ (11/15), but none were NOS^+^ (0/15) (N = 6; Figure 4D-F; table 3). L-Phe (100 mM) activated significantly fewer myenteric neurons (8 ± 1%; n = 323, N = 11; Figure G-I; table 3) compared to mucosal depolarization (P = 0.017; One-way ANOVA, Tukey’s multiple comparisons test; Figure 4J). Of the L-phe responders, most were calbindin^+^ (13/22 neurons), while none were NOS^+^, and many did not label for either marker (9/22 neurons, N = 7). The total proportions of calbindin^+^ and nNOS^+^ neurons examined were consistent across preparations (Table 3). However, the relative proportions of the neurochemical subtypes of responding myenteric neurons differed significantly between stimuli (*P* < 0.0001; χ2 test). A comparison between the cell diameters of responding neurons further highlights the unique populations of responders to each stimulus (Figure 4K-L). Thus, in contrast to nutrient responses in the submucosal plexus, these data indicate that different nutrients activate distinct patterns of myenteric neurons.

Krebs perfusion alone did not evoke any consistent Ca^2+^ responses in the submucosal plexus (n = 93, N = 3) or the myenteric plexus (n = 181 neurons, N = 5), demonstrating that the perfusion pressure applied was not a sufficient mechanical stimulus to evoke mechanosensory responses in our preparations.

### Mucosal nerve endings are not directly sensitive to luminal nutrients

Given that nutrients are transported or passively diffuse across the epithelium, we next tested whether mucosal sensory nerve endings or cell bodies are directly sensitive to nutrients. Glucose (300 mM; Suppl. Movie 7), acetate (100 mM; Suppl. Movie 8), or L-phe (100 mM; Suppl. Movie 9) were applied into a single villus by pressure ejection via a micropipette pushed through the epithelium to directly target the sensory nerve endings. However, this mode of nutrient application did not elicit any responses in myenteric neurons (glucose: n = 175, N = 4; acetate: n = 198, N = 3; L-phe: n = 213, N = 3). The local application of these nutrients by pressure ejection directly onto myenteric ganglia in peeled LMMP preparations also did not elicit specific neuronal responses (glucose: n = 154, N = 3; acetate: n = 181, N = 3; L-phe: n = 271, N = 3). This indicates that mucosal nerve endings and neuronal cell bodies do not directly respond to glucose, acetate, or L-phe that passes across the epithelium, at least at the concentrations tested. Further, this data indicates that the mucosa is first necessary for sensory transduction of the luminal signal to the ENS. To verify the mucosa responds to the nutrient solutions at the selected concentrations, we applied each nutrient to the mucosal surface of wholemount preparations of jejunum from Villin|GCaMP3 mice, which express GCaMP3 in their intestinal epithelium. The nutrient and high K^+^ solutions each elicited defined Ca^2+^ responses in the epithelium (Figure 5).

**Figure 5:**
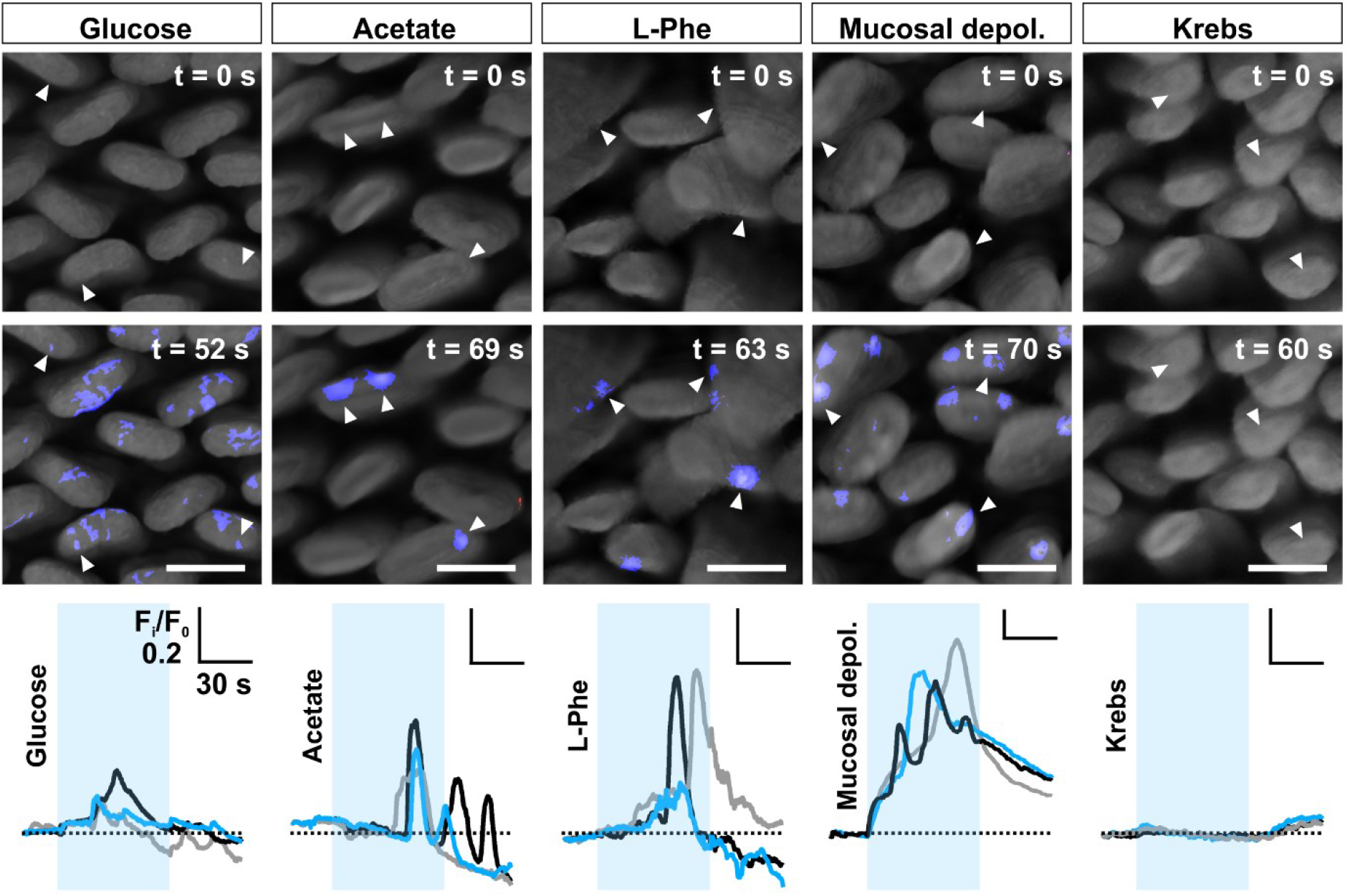
Nutrients elicit defined epithelial Ca^2+^ responses. Representative widefield images of the mucosal surface of Villin | GCaMP3 mouse jejunum during the application of glucose (300 mM), acetate (100 mM), L-phenylalanine (L-Phe, 100 mM), high K^+^ (75 mM) Krebs to depolarise the mucosa (mucosal depol.), or control Krebs buffer, and at baseline (t = 0 s). Regions of villi that displayed an increase in fluorescence intensity in response to each stimulus are highlighted in blue. For each stimulus, examples of Ca^2+^ transients in selected regions of epithelium (as marked by arrowheads) are shown in the bottom panel. Stimuli were applied from t = 20 s to t = 80 s. The timing and duration of stimulus application is indicated by the blue boxes. Scale bars = 200 μm.

### Enteric mucosal nerve endings respond to 5-HT and ATP

To identify potential mucosal mediators involved in transducing luminal signals to the ENS, we assessed their ability to stimulate mucosal nerve endings and activate enteric neurons. Putative mucosal signaling mediators implicated in nutrient sensing were injected into individual villi while imaging Ca^2+^ activity in the underlying myenteric plexus (Figure 6A). The effects of 5-HT, ATP, GLP-1 and CCK-8 were examined (Table 4).

**Figure 6:**
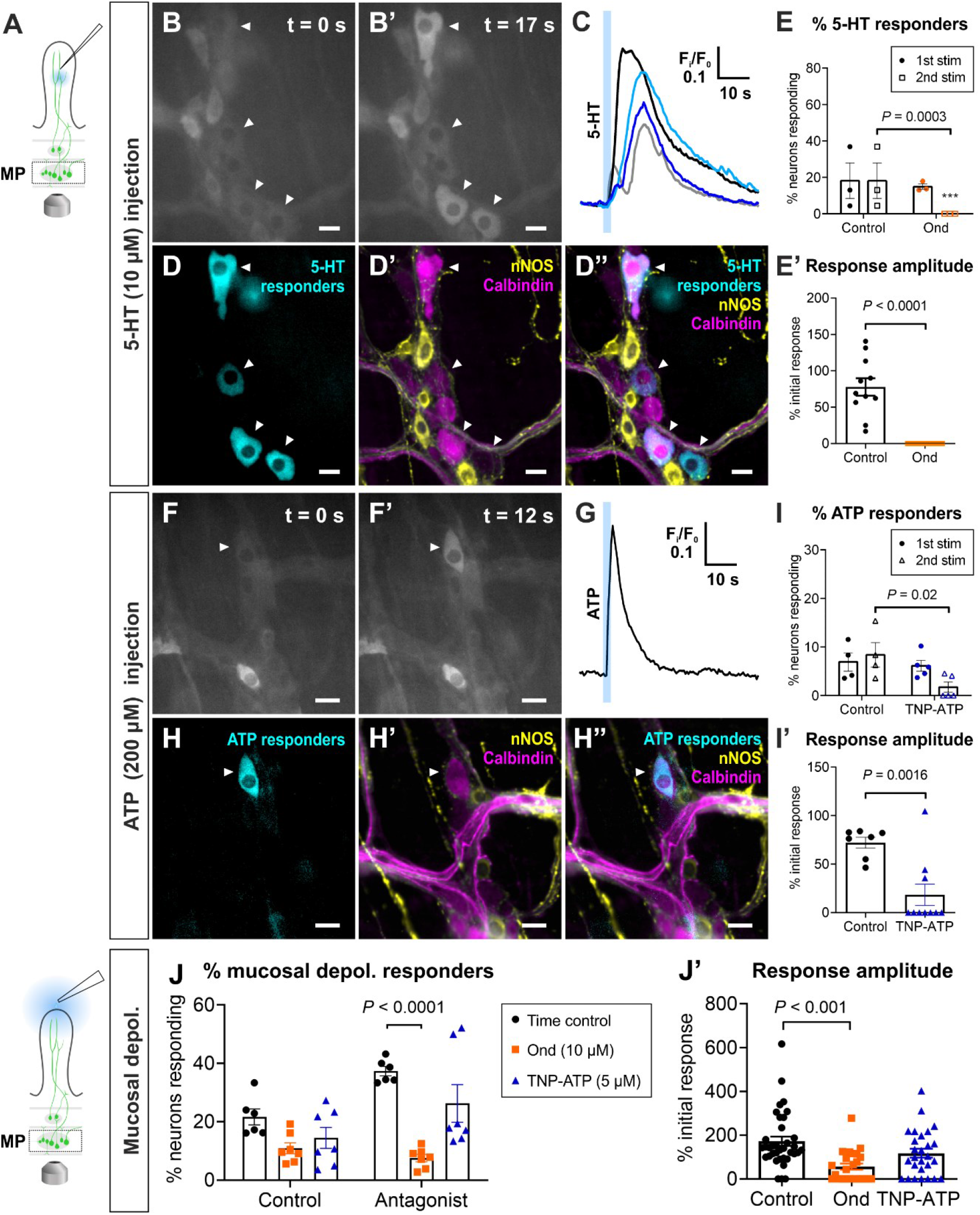
Mucosal nerve endings are sensitive to 5-HT and ATP. **A.** Myenteric neurons were imaged while 5-HT (10 μM) or ATP (200 μM) were injected into a single villus via a micropipette pushed through the epithelium. **B-B’.** 5-HT (10 μM) applied onto the mucosal nerve endings activated a subset of myenteric neurons as marked by white arrowheads. Scale bars = 20 μm. **C.** Corresponding calcium transients in the responding neurons marked in the previous panels. 5-HT was applied t = 10 s. **D.** 5-HT responders are depicted in cyan. **D’.** The corresponding ganglion immunolabelled for calbindin and neuronal nitric oxide synthase (nNOS). **D”.** An overlay of panels **D** and **D’** show that many 5-HT responders are calbindin^+^ neurons. **E-E’.** Two repeated 5-HT injections into a villus evoked reproducible responses in myenteric neurons under control conditions. The 5-HT response was abolished in the presence of the 5-HT_3_ antagonist ondansetron (ond, 10 μM) (Two-way ANOVA, Sidak’s multiple comparisons test). **F-F’.** ATP (200 μM) applied onto the mucosal nerve endings activated myenteric neurons as marked by white arrowhead. Scale bars = 20 μm. **G.** The corresponding calcium transient in the responding neuron. ATP was applied t =10 s. **H.** The ATP responder is depicted in cyan. **H’.** The corresponding ganglion stained for the calbindin and neuronal nitric oxide synthase (nNOS). **H”.** An overlay of panels **H** and **H’** show that the ATP responder was calbindin^+^. **I-I’.** Two repeated ATP injections into a villus evoked reproducible responses in myenteric neurons under control conditions. The ATP response was significantly reduced in terms of the percentage of neurons responding and response amplitudes in the presence of the P2 antagonist TNP-ATP (5 μM). **J.** The percentage of myenteric neurons responding to mucosal depolarization was also significantly inhibited by ondansetron (10 μM), but not TNP-ATP (5 μM), when compared to the corresponding control time point. **J’.** The response amplitudes were also significantly reduced in the presence of ondansetron (10 μM), but not TNP-ATP (5 μM).

GLP-1 (1 μM) (n = 217, N = 4) and CCK-8 (100 nM) (n = 166, N = 3) did not evoke any responses when injected into villi, indicating that these molecules do not act (at least acutely) on enteric mucosal nerve endings. This does not rule out the possibility that these mediators have modulatory roles on enteric nerve activity or that they specifically target extrinsic mucosal nerve endings. Nonetheless, the lack of responses to these stimuli when injected into villi demonstrates that responses elicited were not simply due to mechanical distortion or pressure.

5-HT/5-HT3 signaling is considered a key mucosal sensory pathway in the GI tract involved in transducing chemical and mechanical stimuli (Bertrand, 2009, Bellono et al., 2017, Alcaino et al., 2018). 5-HT (10 μM) injected into the villus evoked reproducible responses in 18 ± 10% myenteric neurons (n = 65, N = 3) (Figure 6B-D”). In accordance with our previous findings (Hao et al., 2020), these responses are mediated via 5-HT3 receptors as they were abolished by ondansetron (10 μM, 10 min; N = 3) (Figure 6E-E’).

There is also evidence for the role of ATP as a sensory mediator in mucosal mechanosensitivity (Cooke et al., 2003). ATP applied to the luminal side of the mucosa was found to activate myenteric AH neurons in guinea pig ileum via P2X receptors (Bertrand and Bornstein, 2002). Here, we demonstrate that ATP (200 μM) applied into a single villus evokes Ca^2+^ responses in 8 ± 2% myenteric neurons (n = 201, N = 4; Suppl. Movie 10); almost all ATP responders were calbindin^+^ (11/12 neurons) but rarely nNOS^+^ (1/12 neurons, N = 4) (Figure 6F-H”). Furthermore, the percentage of ATP responders was significantly reduced to 3 ± 1% by the purinergic P2X receptor antagonist TNP-ATP (5 μM) (n = 148, N = 5; *P* = 0.02, Two-way ANOVA, Sidak’s multiple comparisons test) (Figure 6I-I’). The response amplitude was also inhibited by TNP-ATP (ΔF_i_/F_0_ = 0.03 ± 0.01 vs. time control: ΔΔF_i_/F_0_ = 0.09 ± 0.01; *P* = 0.03, Two-way ANOVA, Sidak’s multiple comparisons test), indicating that the response to ATP is largely mediated by P2X receptors.

### Mucosal depolarization activates serotonergic but not purinergic pathways

Having demonstrated that mucosal nerve endings respond to 5-HT and ATP, we then tested whether there is endogenous release of these mediators upon mucosal depolarization using the 5-HT3 antagonist ondansetron (10 μM) and P2X receptor antagonist TNP-ATP (5 μM). Ondansetron significantly inhibited the proportion of myenteric neurons responding to mucosal depolarization (ondansetron: 8 ± 1% vs. time control: 37 ± 2%; N = 7; *P* = 0.0003, Two-way ANOVA, Sidak’s multiple comparisons test) (Figure 6J). The response amplitude was also substantially reduced (ondansetron: ΔF_i_/F_0_ = 0.06 ± 0.01 vs. time control: ΔF_i_/F_0_ = 0.15 ± 0.01; n = 222; *P* < 0.0001, Two-way ANOVA, Sidak’s multiple comparisons test) (Figure 6J’). Nonetheless, a residual neuronal response was still observed in the presence of ondansetron, suggesting the involvement of signaling mechanisms other than the 5-HT/5-HT3 receptor pathway upon exposure to luminal stimuli. Although mucosal nerve endings were also responsive to ATP, TNP-ATP had no effect on myenteric responses to mucosal depolarization (N = 7; Figure 6J-J’), indicating that ATP acting via P2 receptors is not involved in mediating this response.

### The primary responding neurons to mucosal stimuli are localized to the myenteric plexus

While we have shown that mucosal depolarization activates many calbindin-immunoreactive myenteric neurons, suggesting that they are IPANs, it is unclear whether they respond directly, or indirectly via a synaptic connection in the network. This was addressed using the nicotinic antagonist hexamethonium (hex, 200 μM) to block the major form of excitatory neurotransmission in the ENS. The percentage of myenteric neurons responding prior to and following hex exposure were comparable (24 ± 11% vs. 23 ± 7%, N = 6), demonstrating that the responders are predominantly primary responding neurons (Figure 7B).

**Figure 7:**
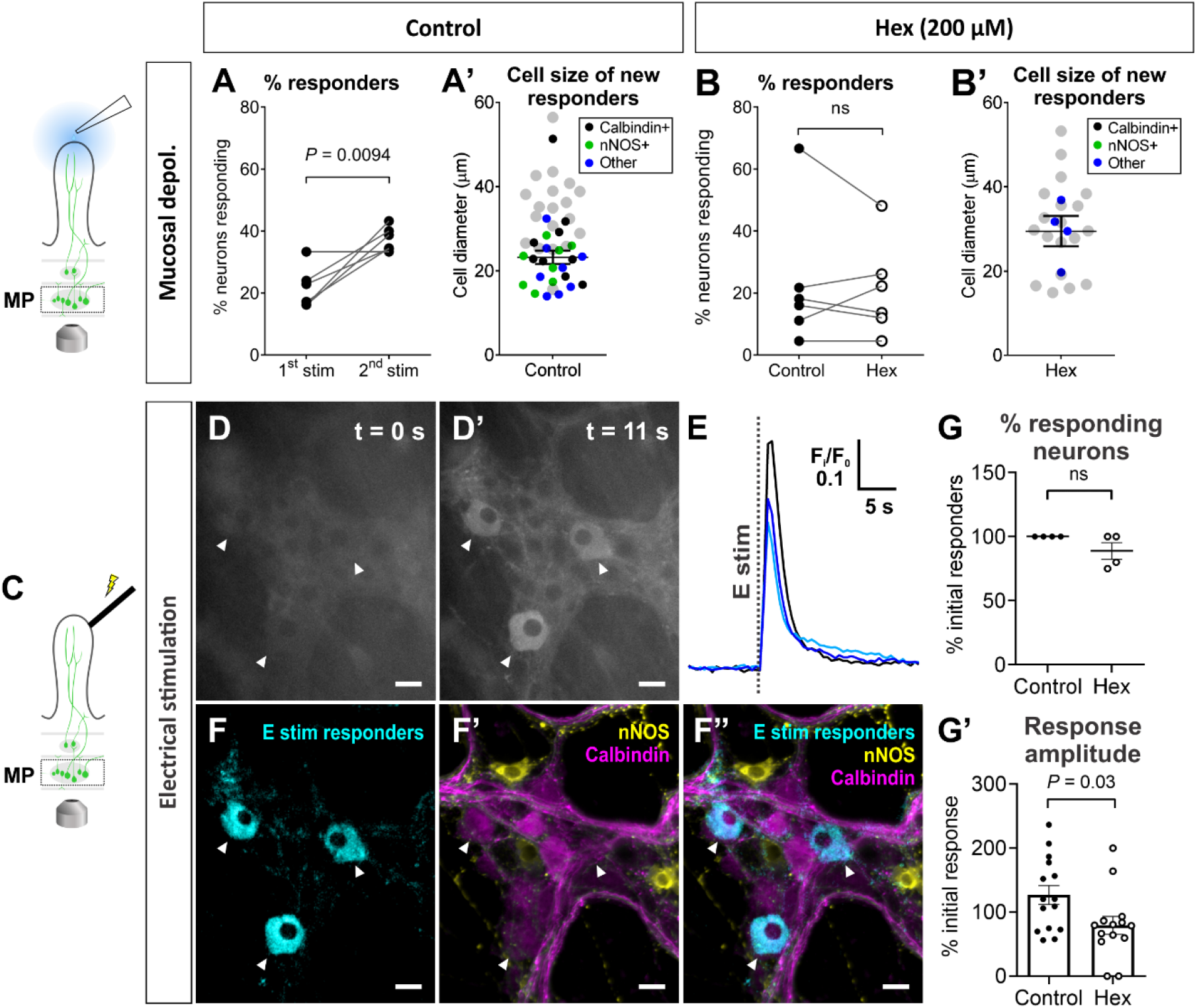
Mucosal depolarization predominantly activates myenteric sensory neurons. **A.** Repeated mucosal depolarization evoked reproducible responses in myenteric neurons, albeit a significant increase in the number of responders with the 2^nd^ stimulation (paired t-test). **A’.** The cell diameter of the new responders were relatively smaller than the initial responders (depicted in grey) and many were nNOS-immunoreactive or did not label for either nNOS or calbindin. **B.** In the presence of hexamethonium (hex; nicotinic antagonist; 200 μM), the percentage of neurons that responded to the 2^nd^ mucosal depolarization was comparable to that of the initial response, demonstrating that the initial responding neurons respond directly to the stimulus. **B’.** Only few new responders were observed in the presence of hex and none of these labelled for nNOS or calbindin. Initial responders are shown in grey. **C.** A focal electrode was placed onto the tip of a villus and stimulated with a train of 20 pulses for 1 s while responses in the myenteric plexus was imaged. **D-D’.** Directly stimulating the mucosal nerve endings electrically evoked responses in a subset of myenteric neurons as marked by the white arrowheads. Scale bars = 20 μm. **E.** Corresponding calcium transients in the responders indicated in the previous panels are shown. The train of pulses was applied at t = 10 s. **F.** Responders to the electrical stimulus are depicted in cyan. **F’.** Immunolabelling of the corresponding ganglion for calbindin and nNOS. **F”.** The overlay of panels **F** and **F’** show that some responders are calbindin^+^. **G-G’.** The percentage of myenteric neurons was not significantly different between controls and in hex (200 μM) conditions, indicating that electrical stimulation of the mucosal nerve endings directly activated myenteric sensory neurons. However, the response amplitudes were significantly reduced in the presence of hex, suggesting that the response is modulated by excitatory synaptic inputs. The neurochemistry of these responders is comparable to those that responded initially to mucosal depolarization.

Interestingly, under control conditions, repeated mucosal depolarization evoked a larger response in the myenteric plexus, with the second stimulation activating significantly more neurons (from 22 ± 3% to 37 ± 2%; n = 175, N = 5; *P* < 0.0094) (Figure 7A). The additional neurons activated by repeated mucosal depolarization were further characterized as mainly neurons with smaller cell bodies (21 ± 2 μm in diameter) that were calbindin^+^ (9/25 neurons), nNOS^+^ (8/25 neurons), or did not label with either marker (Figure 7A’). Taken together, it is likely that these are excitatory and inhibitory motor neurons and/or interneurons that were recruited. The amplified response with the second mucosal depolarization was absent in hex conditions (Figure 7B-B’). This indicates that the additional neurons responding to the repeated mucosal depolarization under control conditions were activated secondarily via nicotinic neurotransmission.

To characterize the myenteric neurons within our field of view that are functionally connected to a single villus, we electrically stimulated the nerve endings using a focal electrode positioned on the villus tip in the presence of hex (Figure 7C). Electrical train stimulation (20 Hz, 1 s) under control conditions elicited reproducible responses in a 12 ± 2% of myenteric neurons (n = 221, N = 4). Similar to the initial mucosal depolarization responders, electrical stimulation predominantly evoked responses in calbindin^+^ neurons (12/15; Figure 7D-F”; Table 4). Hex reduced the amplitude of the response to the 2^nd^ electrical stimulation to 79 ± 15% of the initial response (vs. 135 ± 19% in control conditions), demonstrating that the response is modulated by excitatory synaptic transmission (Figure 7G’). Although not significantly different, the proportion of neurons responding to the 2^nd^ electrical stimulation in the presence of hex was 89 ± 7% of the initial responders, while the same number of neurons responded to both stimulations under control conditions (Figure 7G). This indicates that the myenteric neurons responding to electrical stimulation are predominantly primary responding neurons, with a minor fraction of neurons responding synaptically. Overall, approximately 14 ± 5% of myenteric neurons (n = 107) within our field of view functionally innervate a single villus. This is compatible with the numbers of myenteric neurons responsive to 5-HT (15 ± 3%) applied into a villus, suggesting that the majority (if not all) of the nerve endings supplying the mucosa are sensitive to 5-HT.

Next, we used spinning disk confocal microscopy to examine neuronal responses to mucosal depolarization in both the myenteric and submucosal plexus simultaneously in full thickness preparations (Figure 8; Suppl. Movie 11). The proportion of myenteric and submucosal neurons that responded was 20 ± 5% and 26 ± 14% of total neurons examined, respectively (Figure 8D); these data are consistent with the proportion of responders to mucosal depolarization examined using widefield microscopy (Figures 3-4). Notably, the initial responding neuron was consistently found in the myenteric, rather than the submucosal plexus (MP: n = 111; SMP: n = 39; N = 4). The peak of the Ca^2+^ signal in the initial responding myenteric neuron preceded that of the initial responding submucosal neuron in 3/4 preparations examined (Figure 8A-C). In 1 preparation, only myenteric responses were observed. These data provide further evidence that the primary response to mucosal stimulation is in the myenteric plexus.

**Figure 8:**
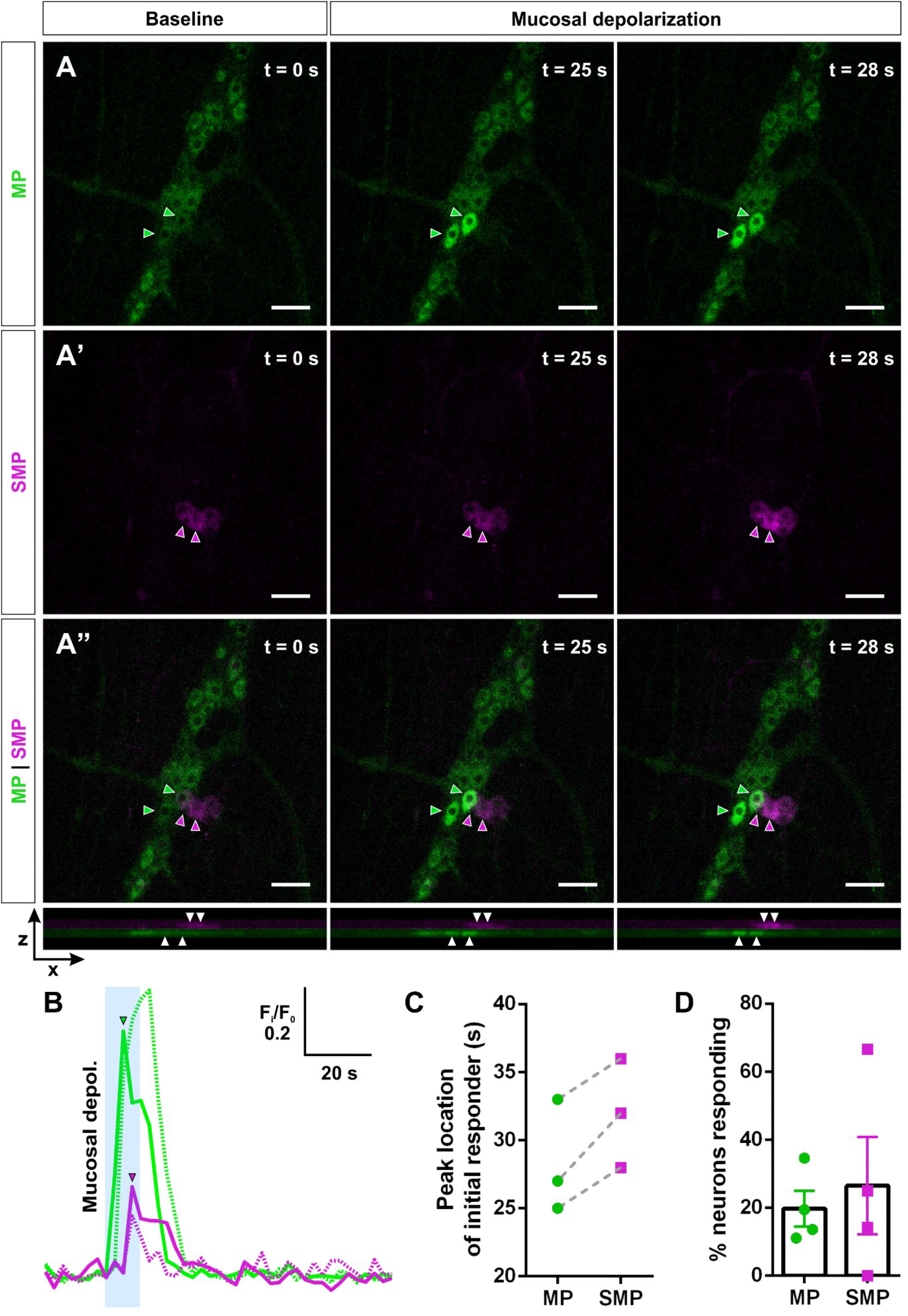
Myenteric neurons respond primarily to mucosal depolarization. **A-A”.** Representative spinning disk confocal microscope recording of Ca^2+^ activity in the myenteric (MP) and submucosal plexus (SMP) in response to mucosal depolarization (from t = 20 s to t = 30 s). Maximum XY projections of the Z stack (25 μm) divided into the **A.** MP and **A’.** SMP. **A”.** Maximum XY and XZ projections of the full stack with the MP and SMP depth colour-coded in green and magenta, respectively. Scale bars = 50 μm. Responding neurons are indicated by color-coded arrowheads. **B.** Corresponding traces of Ca^2+^ responses in the myenteric (green) and submucosal neurons (magenta) indicated in panels **A-A”**, depicted as a change in fluorescence over time. Arrowheads mark the Ca^2+^ peak of the initial responders. **C.** The Ca^2+^ peak of the initial responding myenteric neuron consistently precedes that of the initial responding submucosal neuron in corresponding preparations (N = 3). **D.** The proportion of myenteric and submucosal neurons that responded to mucosal depolarization.

### Submucosal responses to mucosal stimulation are predominantly synaptically-mediated

To determine whether there are also IPANs in the submucosal plexus that respond directly the mucosal depolarization, we also examined submucosal responses in the presence of hex. In contrast with myenteric neurons, the number of submucosal neurons responding to mucosal depolarization were significantly inhibited by hex (control: 27 ± 6% vs. hex: 13 ± 4%, N = 8) (Figure 9B). The response amplitude was also significantly reduced in hex conditions (ΔF_i_/F_0_ = 0.10 ± 0.01 vs. ΔF_i_/F_0_ = 0.02 ± 0.01 in control conditions, n = 162) (Figure 9B’). Submucosal responses to repeated mucosal depolarization under control conditions evoked comparable responses in terms of numbers responding (1^st^: 27 ± 6% vs. 2^nd^: 22 ± 5%; N = 7) and response amplitudes (1^st^: ΔF_i_/F_0_ = 0.08 ± 0.01 vs. 2^nd^: ΔF_i_/F_0_ = 0.08 ± 0.01; n = 132) (Figure 9A-A’). Collectively, these data show that submucosal responses are largely activated indirectly via nicotinic transmission.

**Figure 9:**
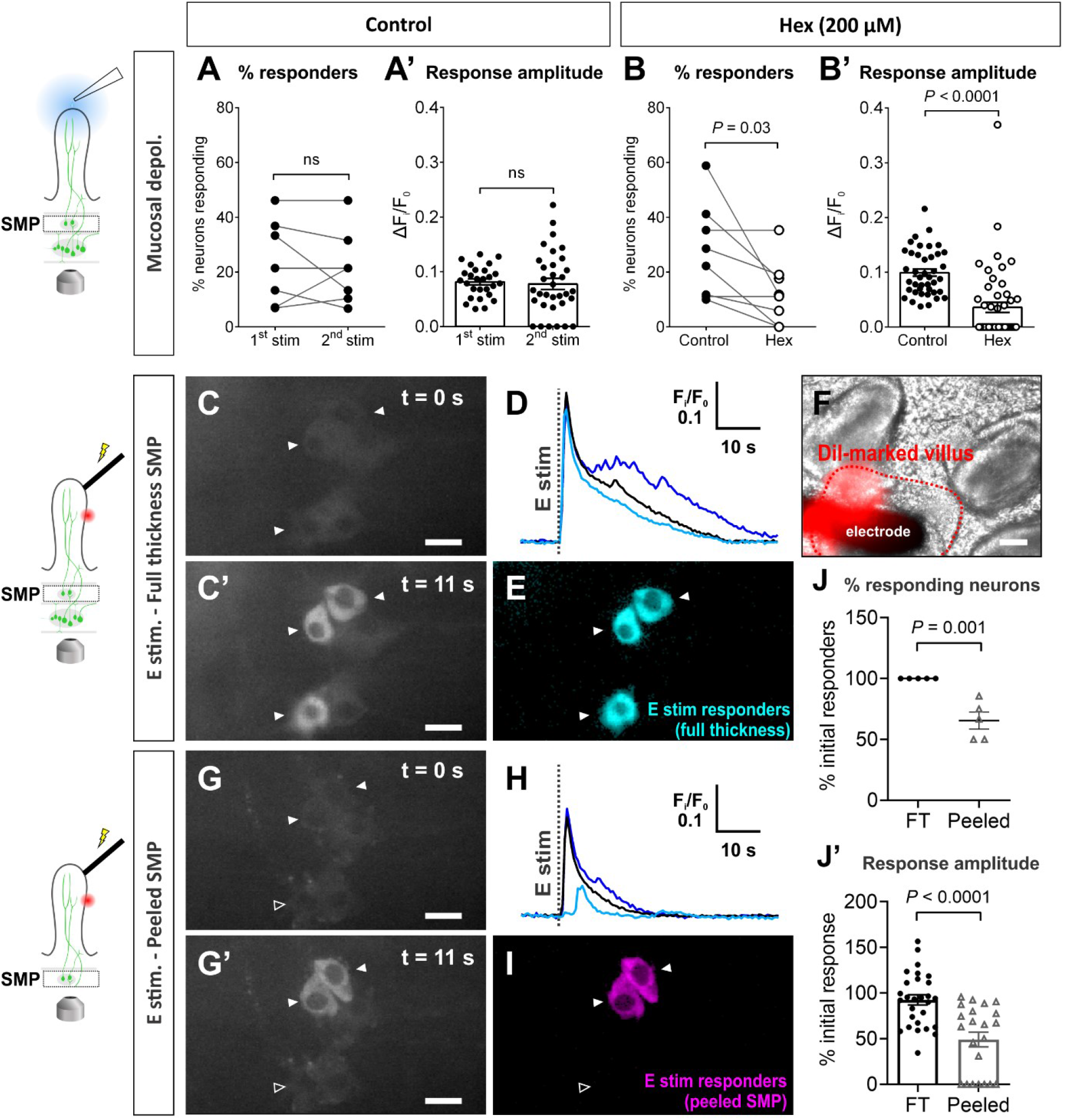
Submucosal responses to mucosal stimulation are mediated by nicotinic transmission and via the myenteric plexus. **A-A’.** Repeated mucosal depolarization evoked reproducible responses in submucosal neurons under control conditions. **B-B’.** Submucosal responses to mucosal depolarization were significantly inhibited in the presence of hexamethonium (hex; nicotinic antagonist, 200 μM). **C-C’.** Submucosal neurons activated by focal stimulation of a villus (20 Hz, 1 s) in a full thickness preparation. Responders are indicated by white arrowheads. Scale bars = 20 μm. **D.** Corresponding calcium transients in the responding neurons. The train of pulses was applied at t = 10 s. **E.** Responders are shown in cyan. **F.** The villus stimulated is marked with a red fluorescent dye (DiI) for the purpose of relocating it after peeling the preparation. **G.** The same villus marked by DiI was stimulated again following the removal of the myenteric plexus by microdissection. Only a subset of the initial responders were activated. **H.** Corresponding calcium transients in marked neurons are shown. A reduction in the amplitude in apparent in the remaining responders. **I.** Responders to the electrical stimulus in the peeled preparation are shown in magenta. **J.** There is a significant reduction in percentage of initial submucosal neurons responding following the removal of the myenteric plexus. **J’.** The response amplitudes of the responders are also significantly reduced in the peeled vs. full thickness (FT) preparation.

Given that the primary responders to mucosal depolarization are predominantly situated in the myenteric plexus and submucosal responses largely occur synaptically, we examined the extent to which submucosal ganglia can receive synaptic inputs from the myenteric plexus. Responses to electrical stimulation of mucosal nerve endings in a single villus in the presence and absence of the myenteric plexus were compared. First, a train of 20 pulses (20 Hz, 1 s) was applied to the villus via a focal electrode in full thickness preparations. This evoked Ca^2+^ transients in 31 ± 5% of submucosal neurons (n = 73, N = 5) (Figure 9C-E). The same villus (marked by DiI applied on the epithelium) was electrically stimulated again after peeling away the myenteric plexus (Figure 9F). Removing the myenteric plexus reduced the number of initial submucosal responders by 35 ± 7% (n = 73, N = 5) and the response amplitude was reduced to 49 ± 8% of the initial response (Figure 9G-J’). Repeated electrical stimulation evoked reproducible responses in full thickness preparations, where the initially responding neurons were all activated by the 2^nd^ stimulation and their response amplitude was 93 ± 6% of the initial response (N = 5; Figure 9J-J’). Thus, these data show that a subset of submucosal neurons is activated only secondarily via a myenteric neuron intermediate.

## Discussion

In this study, we investigated the communication of luminal information, with a focus on nutrient signals, to the ENS in the intact gut wall. Our results show that the first order neurons (i.e. IPANs) that respond to luminal stimuli are situated in the myenteric plexus, and that different patterns of myenteric neurons are activated in response to different nutrients. The sensory nerve endings and myenteric cell bodies did not respond directly to the nutrients indicating that they are first sensed by the epithelium and intermediate mucosal mediators are necessary to signal to the ENS. Consistent with previous work (Bertrand, 2009, Bellono et al., 2017, Alcaino et al., 2018, Hao et al., 2020), we confirm – at the level of a single villus – that 5-HT acting via 5-HT3 receptors is one of the key signaling processes involved in communicating the luminal information from EECs. Interestingly, the primary response to mucosal stimuli was predominantly localized to the myenteric plexus, rather than the submucosal plexus, despite the latter being physically closer to the epithelium. Following the activation of myenteric IPANs, the information is then transmitted to cholinergic submucosal neurons via nicotinic transmission.

### The ENS discriminates between different luminal signals

In order for the ENS to generate the appropriate motor and secretory behaviors, it is crucial for it to first decipher the nature of its luminal contents. Different motor patterns are observed *in vivo* when fed a nutrient vs. a non-caloric meal, where the nutrient meal produced mixing motor patterns to facilitate digestion and absorption, the non-caloric meal predominantly evoked propulsive contractions (Schemann and Ehrlein, 1986). Or in the case when presented with the diarrhoea-inducing cholera toxin together with a nutritious fatty acid, the intestine apparently overrides the signal to trigger the mixing motor patterns, and propulsive contractions persist (Fung et al., 2010), presumably prioritizing the elimination of the noxious agent. However, the extent to which the gut discriminates between different substances within the luminal contents is unclear. In our current study, we demonstrate that the ENS is capable of distinguishing between a luminal sugar, fatty acid, and amino acid signal. The different types of nutrients applied to the mucosa elicited differential patterns of myenteric neuronal activation, reflected by the different populations of responders as identified by their neurochemical subtypes and size of their cell bodies. On the other hand, all responders in the submucosal plexus were cholinergic neurons. This suggests that while different nutrient signals trigger the activation of different local sensory and motor circuits, they ultimately converge onto common secretomotor and/or vasodilator pathways in the submucosal plexus. Further, the full breadth of neuronal activation likely extends beyond our field of view, particularly with respect to ascending and descending interneurons and motor neurons.

### 5-HT mediates communication between EECs and enteric nerves via 5-HT3 receptors

Our experiments using Villin|GCaMP3 tissues showed that the different nutrient solutions tested evoked distinct Ca^2+^ responses in the mucosal epithelium, while mucosal nerve endings and neuronal cell bodies were not directly sensitive to these nutrients. This is compatible with the nutrient signal being first detected at the level of the epithelium i.e. by EECs which contain various signaling mediators such as 5-HT (Diwakarla et al., 2017, Fothergill and Furness, 2018). Blocking 5-HT3 receptors significantly inhibited the response to mucosal depolarization, supporting the notion that upon depolarization of EECs, mucosal 5-HT is released to activate intrinsic sensory nerve endings (Bertrand et al., 2000). While the response can also be mediated by neuronal 5-HT within the circuitry, it has been shown that only a small subset of myenteric S neurons exhibit 5-HT3-mediated fast EPSPs (at least in guinea pig ileum) (Zhou and Galligan, 1999). Furthermore, our data shows that the myenteric neurons responding to mucosal depolarization are predominantly calbindin^+^ IPANs. The data also indicates the involvement of signaling mediators other than 5-HT and/or other 5-HT receptors as the response was not abolished by ondansetron.

ATP was also considered as a candidate molecule linking EECs to the activation of myenteric IPANs. While it effectively stimulated a P2X-mediated response in myenteric neurons when applied to mucosal nerve endings, the myenteric response to mucosal depolarization was P2X-insensitive, suggesting that ATP is not a major mediator released by depolarizing EECs. Nevertheless, ATP is not released solely from EC cells but also from epithelial cells upon mechanical stimulation of the mucosa (Cooke et al., 2003). Thus, it is feasible that ATP plays a role in mechanosensory transduction.

GLP-1 is released from EECs following food intake and in addition to its incretin activity, it also regulates intestinal motility (Amato et al., 2010). GLP-1 receptor-expressing nerve fibers are present in the mucosa (Richards et al., 2014), but whether these are of intrinsic or extrinsic origin, or both, was unclear. CCK is another intestinal hormone secreted from EECs following a meal. While both molecules are thought to be mediators that can signal to IPANs, our data does not support the involvement of mucosal release of GLP-1 or CCK-8 in the activation of enteric sensory nerve endings. Further examination of the activation of enteric IPANs by mucosal mediators will be informed by single cell RNA-sequencing (scRNA-seq) data on EEC-subtype enriched genes encoding for key signaling molecules (Haber et al., 2017), together with data on the gene expression of corresponding receptors in putative IPANs (Zeisel et al., 2018, Drokhlyansky et al., 2020).

### Some submucosal neurons receive luminal information via myenteric neurons

With the various modes of mucosal stimulation including depolarization, nutrient perfusion, electrical stimulation of mucosal nerve endings, and injection of 5-HT or ATP into a villus, the most prominent responders are calbindin-immunoreactive myenteric neurons with larger cell bodies than non-responders (Qu et al., 2008). This is consistent with the initial responders of the luminal signal being myenteric IPANs (Dogiel type II neurons) (Furness et al., 2004). Until recently, the presence of conventional IPANs in the submucosal plexus has been elusive (Mao et al., 2006, Wong et al., 2008, Mongardi Fantaguzzi et al., 2009, Foong et al., 2014). Latest work shows that there are IPANs present in the mouse SMP based on neuronal morphology, projection patterns, and advillin expression (Melo et al., 2020), as well as scRNA-seq data (Drokhlyansky et al., 2020). Hence, submucosal neurons can receive luminal information directly and/or secondarily via neurons residing in the myenteric plexus. We found that in the myenteric plexus, neurons were primarily first-order responders to mucosal depolarization, while a substantial proportion of the response in cholinergic submucosal neurons was synaptically-mediated via nicotinic transmission (Figure 10). Furthermore, our data show that upon stimulating mucosal nerve endings, many submucosal neurons receive synaptic inputs via the myenteric plexus. These findings are compatible with previous work in guinea pig ileum demonstrating that many submucosal S neurons receive nicotinic inputs from myenteric neurons following electrical stimulation of the myenteric plexus (Moore and Vanner, 2000). Additionally, a component of the submucosal response to mucosal stimulation was insensitive to synaptic blockade and independent of the myenteric plexus. These findings provide support of the activation of putative submucosal IPANs. It is unlikely that submucosal neurons were activated antidromically as both cholinergic and VIPergic secretomotor neurons which supply the mucosa are abundant, while submucosal responders were almost exclusively cholinergic (Mongardi Fantaguzzi et al., 2009).

**Figure 10.**
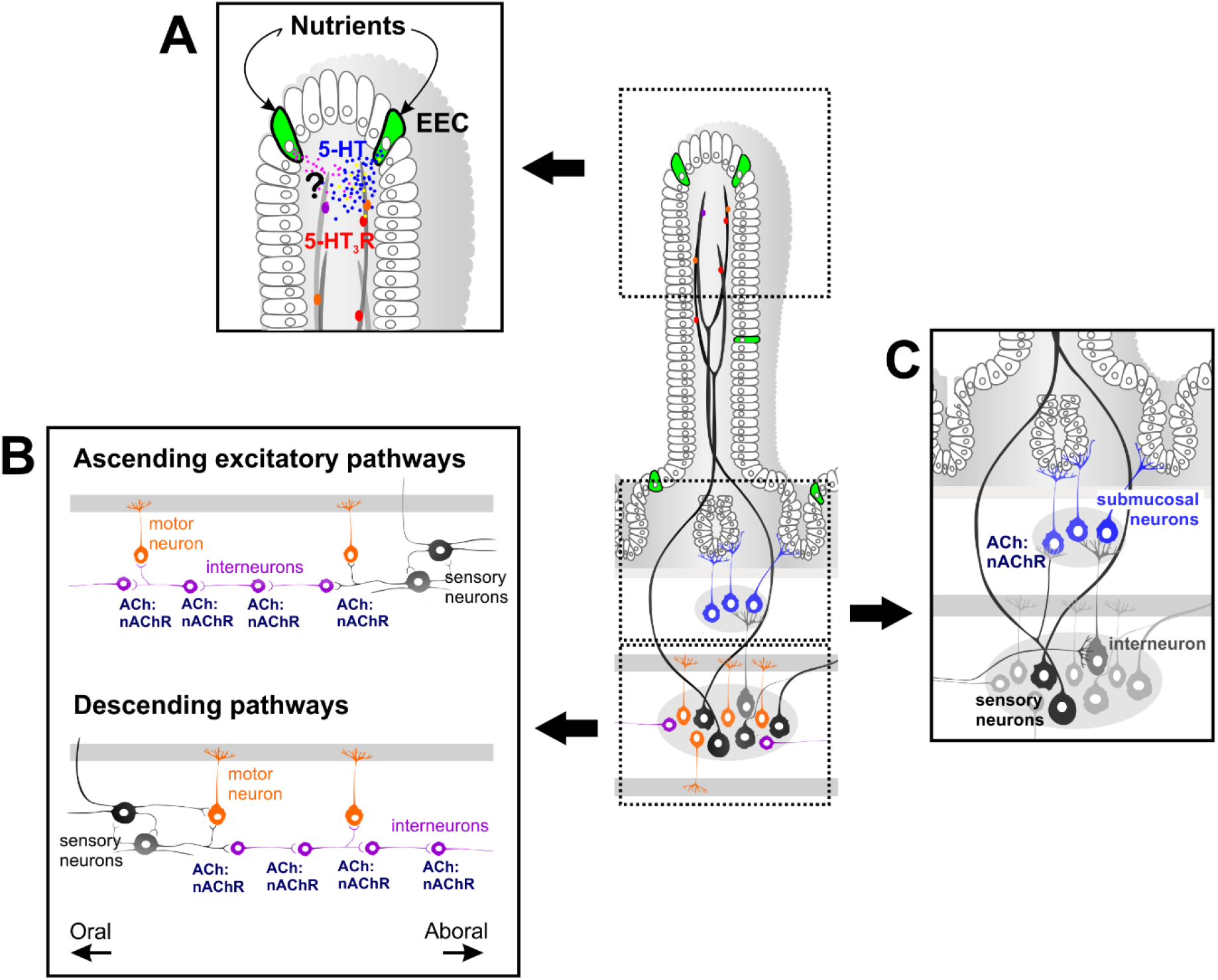
Simplified schematic of the proposed model for the detection of luminal nutrient information by the murine small intestine ENS. **A.** Luminal nutrients are first sensed at the mucosal epithelium via specialised ‘sensor’ cells, with enteroendocrine (EEC) cells, including enterochromaffin cells which release 5-HT upon activation. The basolateral release of 5-HT then stimulates 5-HT_3_ receptor (5-HT_3_R)-expressing intrinsic sensory nerve endings that supply the villi. Other mucosal mediators that may activate intrinsic nerve endings and their corresponding neuronal receptors are yet to be established. **B.** Intrinsic sensory neurons (IPANs) that project to the mucosa predominantly reside in the myenteric plexus and respond primarily to the luminal stimulus. Interneurons and motor neurons of ascending excitatory and/or inhibitory descending pathways may be activated secondarily via nicotinic transmission. **C.** Some submucosal neurons respond secondarily via interneurons and/or myenteric sensory neurons, and their responses are mediated by nicotinic transmission.

### Conclusions

Taken together, our study shows that luminal information from the mucosa is first transmitted to the myenteric plexus. Sensory transduction of luminal signals involves (but is not limited to) 5-HT release, which subsequently activates, via the 5-HT3 receptor, myenteric IPANs. Myenteric interneurons and motor neurons, as well as submucosal neurons are activated secondarily via nicotinic transmission. Notably, specific nutrients differentially activated specific patterns of myenteric, but not submucosal neurons. Thus, while nutrient signals can trigger the activation of different local sensory and motor circuits, they ultimately converge onto common submucosal pathways. Future work will be necessary to determine the full extent of neuronal activation beyond the local site of stimulation examined in this study, particularly regarding ascending and descending interneurons and associated motor neurons.

## Materials and Methods

### Mice and tissue preparation

Adult *Wnt1-Cre;R26R-GCaMP3* and *Villin-CreERT2;R26R-GCaMP3* mice of either sex were killed by cervical dislocation. All procedures are approved by the Animal Ethics Committee of the University of Leuven (Belgium). *Wnt1-Cre;R26R-GCaMP3* mice express the fluorescent Ca^2+^ indicator GCaMP3 in all enteric neurons and glia, while GCaMP3 is expressed in the gut epithelium in *Villin-CreERT2;R26R-GCaMP3* mice (Danielian et al., 1998, el Marjou et al., 2004, Zariwala et al., 2012).

A 5 cm piece of jejunum was collected and immersed in Krebs solution (in mM: 120.9 NaCl, 5.9 KCl, 1.2 MgCl_2_, 1.2 NaH_2_PO_4_, 14.4 NaHCO_3_, 11.5 glucose, and 2.5 CaCl_2_) bubbled with 95% oxygen/5% carbon dioxide. The jejunal segment was opened by cutting along the mesenteric border and pinned flat with the mucosal side up in a silicone elastomer-coated dish. The intestinal contents were removed with Krebs washes and any remnants were carefully removed using forceps. Experiments were limited to a maximum of 3 hours following tissue collection to minimize potential effects of compromised mucosal integrity.

For Ca^2+^ imaging of neuronal activity, either full thickness preparations of *Wnt1-Cre;R26R-GCaMP3* tissues with all the layers of the intestinal wall intact or microdissected submucosal preparations with the mucosa intact, were stretched across an inox ring with the plexus facing up and stabilized with an outer rubber O-ring (Vanden Berghe et al., 2002). Up to 4 ring preparations were obtained from each jejunal segment. In some experiments, the connections between the myenteric plexus and mucosa were severed by peeling apart the gut layers between the submucosal plexus and circular muscle. The two separate layers were then positioned back in place and mounted between the rings.

### Live Ca^2+^ imaging of epithelial activity

*Villin-CreERT2;R26R-GCaMP3* tissues that were stretched and pinned flat with the mucosal side up in a silicone elastomer-coated dish were continuously superfused with Krebs solution bubbled with 95% O_2_|5% CO_2_ throughout the experiment and imaged at room temperature (RT). Imaging was performed using a Leica M165 FC fluorescent stereomicroscope (Leica), fitted with an X-Cite 200DC Stabilized Fluorescence Light Source (Lumen Dynamics) and ORCA-Flash4.0 V3 Digital CMOS camera (Hamamatsu). The mucosa was perfused with stimulating solutions using a micropipette tip positioned at the edge of the imaged field of view. Perfusion was achieved through a gravity-assisted tubing system. Recordings were acquired at 2 Hz with a duration of 2 min, and 5 min between each stimulus application.

### Live Ca^2+^ imaging of neuronal activity

Ring preparations were kept at room temperature in glass coverslip-bottom chambers and constantly perfused with Krebs solution bubbled with 95% oxygen/5% carbon dioxide. Live Ca^2+^ imaging of Wnt1|GCaMP3 ring preparations was performed on either a widefield or a spinning disk confocal microscopy setup. Some preparations were imaged using a 20x objective on a Zeiss Axiovert 200M microscope, equipped with a monochromator (Poly V) and a cooled CCD camera (Imago QE) (TILL Photonics). Images were acquired using TILLVISION software (TILL Photonics). A separate set of experiments were conducted using an inverted spinning disk confocal microscope (Nikon Ti – Andor Revolution – Yokogawa CSU-X1 Spinning Disk [Andor, Belfast, UK]) equipped with a Nikon 20× lens (NA 0.8). GCaMP3 was excited at 488 nm and 3-dimensional (3D) stacks (25 - 29 μm) were acquired using a Piezo Z Stage controller.

#### Mucosal application of nutrient stimuli

Glucose (300 mM), acetate (100mM), or L-phenylalanine (L-phe, 100 mM) were locally applied for 30 s onto the mucosal surface via a perfusion pipette positioned above the imaged myenteric or submucosal plexus. High K^+^ (75 mM) Krebs solution was applied onto the mucosa for 10 s to broadly depolarize EECs and EC cells which are electrically excitable. For some experiments, tissues were perfused with various antagonists for 10 min prior to the second mucosal stimulus.

#### Spritz application of agonist into a single villus

To mimic the basolateral release of mucosal mediators, 5-HT, ATP, CCK-8, or GLP-1, were applied by pressure ejection (Picospritzer II;10 psi, 2 sec, General Valve cooperation) using a micropipette (tip diameter of 10-20 μm) impaled through the epithelial layer and into a single villus as previously described (Hao et al., 2020).

#### Electrical stimulation

To stimulate the nerve endings contained within a single villus, a train of 300 μs pulses (20 Hz, 1 s, 30V) was applied using a focal electrode (50 μm diameter tungsten wire) gently lowered onto the villus tip, coupled to a Grass stimulation unit. The lipophilic tracer DiI (dissolved in EtOH) was applied onto the epithelium of villi by pressure ejection (Picospritzer II; 10 psi, 1 s) as a means of marking and relocating a single villus.

### Nutrients and drugs

Glucose (300 mM; Millipore) was prepared in distilled water. Acetate (100 mM) was prepared with acetic acid (100 mM; Emsure) in a modified Krebs solution (in which the concentration of NaCl was reduced to 20 mM) and neutralized using NaOH (1 M). L-phenylalanine (L-phe; 100 mM; Sigma) was also prepared in a modified Krebs solution with the concentration of NaCl adjusted to maintain the osmolarity. The pH of the nutrient solutions was adjusted to 7.4. 5-HT (Sigma). The concentrations of nutrient solutions were selected to be within the range that is typically ingested. For instance, popular sweetened beverages contain around 300 mM glucose (Ventura et al., 2011), vinegar can contain up to 1.17 M acetic acid (Akiba et al., 2015), and the concentration of L-phe in beef is around 70 mmol/kg (Reitelseder et al., 2020).

ATP (Tocris), GLP-1 (7-36)-amide (Phoenix Pharmaceuticals), CCK-8 (PolyPeptides Laboratories), hexamethonium bromide (Sigma), ondansetron (Sigma), and TNP-ATP (Tocris) were prepared as stock solutions and diluted to the final concentration using standard Krebs solution.

### Analysis

Ca^2+^-imaging analyses were performed with custom-written routines in Igor Pro (Wavemetrics, Lake Oswego, Oregon, USA) (Li et al., 2019). Regions of interest were drawn over each cell to calculate the average [Ca^2+^]_i_ signal intensity. Values were then normalized to the baseline fluorescence intensity (F_i_/F_0_). Responses were considered when the [Ca^2+^]_i_ signal increased above baseline by at least 5 times the intrinsic noise. [Ca^2+^]_i_ peaks were calculated for each response, with the peak amplitude taken as the maximum increase in [Ca^2+^]_i_ from baseline (ΔF_i_/F_0_. Image overlays highlighting active regions of Ca^2+^ activity in *Villin-CreERT2;R26R-GCaMP3* tissues were created in Fiji using the AQuA plugin (Wang et al., 2019).

### Post-hoc immunohistochemistry

Following Ca^2+^ imaging experiments, some full thickness ring preparations were fixed in 4% formaldehyde/PBS for 90 min at room temperature. Fixed tissues were first separated into submucosal and myenteric plexus preparations by microdissection. The submucosal plexus was isolated by dissecting the submucosa and mucosa free from the underlying muscle and myenteric layers, and the mucosa was then gently scraped off. Myenteric plexus preparations were obtained by removing the circular muscle from the remaining preparation.

Submucosal and myenteric plexus preparations were then processed for immunohistochemistry. Tissues were incubated first in a blocking solution for 2 h at room temperature (4% donkey serum/0.5% triton X-100) and then in primary antisera (Table 1) for 48-72 h at 4°C. Tissues were washed in PBS (3 x 10 min) before incubating in secondary antisera (Table 1) for 2 h at room temperature, and then washed again in PBS (3 x 10 min) prior to mounting onto slides using Citifluor (Citifluor Ltd., Leicester, UK).

**Table 1.**
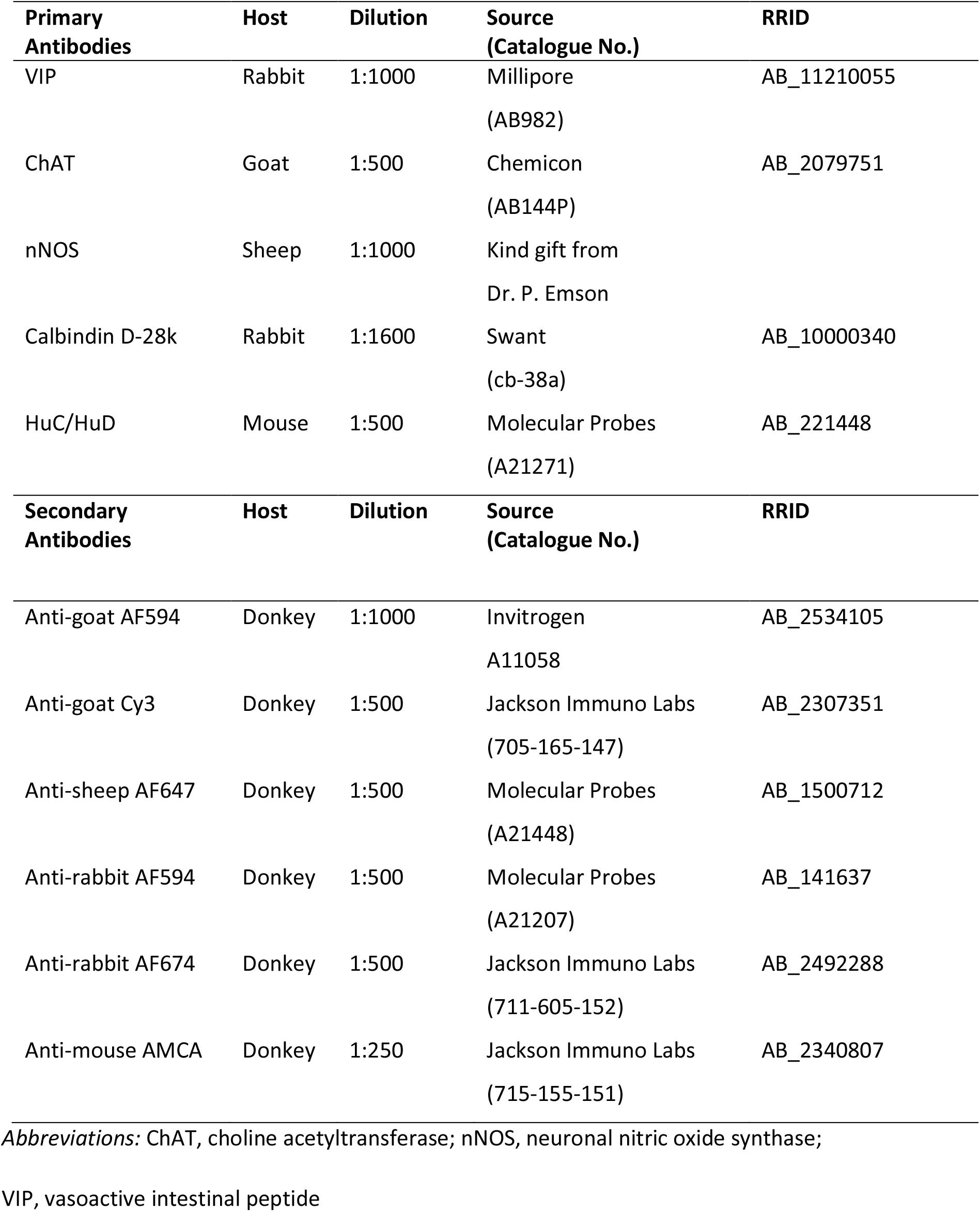
Primary and secondary antibodies used for immunohistochemistry.

| **Primary Antibodies** | **Host** | **Dilution** | **Source (Catalogue No.)** | **RRID** |
| --- | --- | --- | --- | --- |
| VIP | Rabbit | 1:1000 | Millipore<br>(AB982) | AB_11210055 |
| ChAT | Goat | 1:500 | Chemicon<br>(AB144P) | AB_2079751 |
| nNOS | Sheep | 1:1000 | Kind gift from<br>Dr. P. Emson |  |
| Calbindin D-28k | Rabbit | 1:1600 | Swant<br>(cb-38a) | AB_10000340 |
| HuC/HuD | Mouse | 1:500 | Molecular Probes<br>(A21271) | AB_221448 |
| **Secondary Antibodies** | **Host** | **Dilution** | **Source (Catalogue No.)** | **RRID** |
| Anti-goat AF594 | Donkey | 1:1000 | Invitrogen<br>A11058 | AB_2534105 |
| Anti-goat Cy3 | Donkey | 1:500 | Jackson Immuno Labs<br>(705-165-147) | AB_2307351 |
| Anti-sheep AF647 | Donkey | 1:500 | Molecular Probes<br>(A21448) | AB_1500712 |
| Anti-rabbit AF594 | Donkey | 1:500 | Molecular Probes<br>(A21207) | AB_141637 |
| Anti-rabbit AF674 | Donkey | 1:500 | Jackson Immuno Labs<br>(711-605-152) | AB_2492288 |
| Anti-mouse AMCA | Donkey | 1:250 | Jackson Immuno Labs<br>(715-155-151) | AB_2340807 |

*Abbreviations:* ChAT, choline acetyltransferase; nNOS, neuronal nitric oxide synthase;
VIP, vasoactive intestinal peptide

**Table 2.** Submucosal responses to various stimuli perfused onto the mucosa. Mean ± SEM of the percentage of neurons examined within the field of view (n = total number of neurons examined).

|  | % responders | % responders that are ChAT<sup>+</sup> | % total ChAT<sup>+</sup> neurons examined | % responders that are VIP<sup>+</sup> | % total VIP<sup>+</sup> neurons examined |
| --- | --- | --- | --- | --- | --- |
| **Mucosal depolarization** | 24 ± 6%<br>(n = 132) | 96 ± 4%<br>(n = 76) | 55 ± 4%<br>(n = 76) | 4 ± 4%<br>(n = 76) | 45 ± 4%<br>(n = 76) |
| **Glucose** | 21 ± 4%<br>(n = 106) | 100%<br>(n = 77) | 52 ± 5%<br>(n = 77) | 0%<br>(n = 95) | 40 ± 5%<br>(n = 95) |
| **Acetate** | 19 ± 6%<br>(n = 84) | 100%<br>(n = 63) | 66 ± 5%<br>(n = 63) | 0%<br>(n = 63) | 34 ± 5%<br>(n = 63) |
| **L-phe** | 19 ± 5%<br>(n = 195) | 96 ± 4%<br>(n = 126) | 59 ± 3%<br>(n = 126) | 4 ± 4%<br>(n = 126) | 41 ± 3%<br>(n = 126) |
| **Krebs** | 0<br>(n = 83) |  |  |  |  |

**Table 3.** Myenteric responses to various stimuli perfused onto the mucosa. Mean ± SEM of the percentage of neurons examined within the field of view (n = total number of neurons examined).

|  | % responders | % responders that are calbindin<sup>+</sup> | % total calbindin<sup>+</sup> neurons examined | % responders that are nNOS<sup>+</sup> | % total nNOS<sup>+</sup> neurons examined |
| --- | --- | --- | --- | --- | --- |
| **Mucosal depolarization** | 22 ± 3%<br>(n = 175) | 81 ± 8%<br>(n = 115) | 56 ± 5%<br>(n = 115) | 9 ± 5%<br>(n = 115) | 20 ± 4%<br>(n = 115) |
| **Glucose** | 17 ± 6%<br>(n = 183) | 59 ± 15%<br>(n = 183) | 45 ± 9%<br>(n = 183) | 19 ± 9%<br>(n = 183) | 19 ± 3%<br>(n = 183) |
| **Acetate** | 12 ± 2%<br>(n = 153) | 80 ± 9%<br>(n = 153) | 50 ± 7%<br>(n = 153) | 0%<br>(n = 153) | 27 ± 4%<br>(n = 153) |
| **L-phe** | 9 ± 2%<br>(n = 343) | 58 ± 12%<br>(n = 343) | 33 ± 4%<br>(n = 343) | 0%<br>(n = 343) | 24 ± 2%<br>(n = 343) |
| **Krebs** | 0.5 ± 0.5%<br>(n = 1/181) | | | | |

**Table 4.**
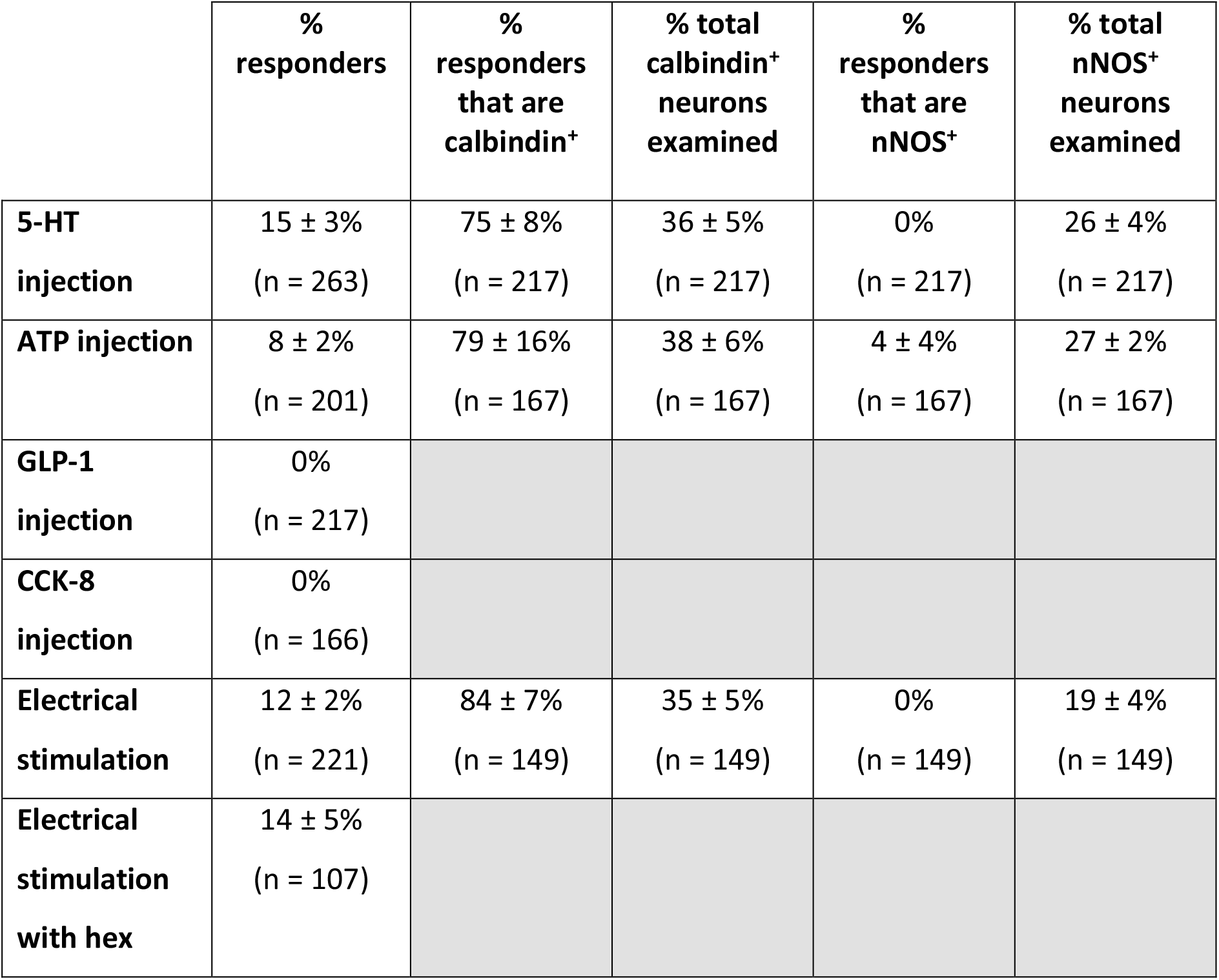
Myenteric responses to various stimuli applied to individual villi. Mean ± SEM of the percentage of neurons examined within the field of view (n = total number of neurons examined).

Immunostained preparations were viewed under an epifluorescence microscope (BX 41 Olympus, Olympus, Aartselaar, BE). Images were acquired with an XM10 (Olympus) camera using Photoshop software, and were adjusted for contrast and brightness before overlay and quantification. Confocal images were recorded using a Zeiss LSM780 microscope (Zeiss).

### Data and statistics

Data are presented as mean ± the standard error of the mean (SEM), with ‘n’ = number of neurons and ‘N’ = number of preparations examined. At least 3 mice were used for each set of experiments. One-way analysis of variance (ANOVA) followed by Dunnett’s *post hoc* test (where appropriate) were conducted to determine statistical significance, unless specified otherwise. *P* < 0.05 was considered significant. Analyses were performed using GraphPad Prism 8.0 (GraphPad softwares, San Diego, California, USA).

## Supporting information

Suppl. Movie 10 - MP activation with intravillus ATP

Suppl. movie 2 - SMP activation by mucosal glucose

Suppl. movie 2 - MP activation by mucosal glucose

Suppl. movie 3 - SMP activation by mucosal acetate

Suppl. movie 4 - MP activation by mucosal acetate

Suppl. movie 5 - SMP activation by mucosal L-phe

Suppl. movie 6 - MP activation by mucosal L-phe

Suppl. movie 7 - villus epithelium activation by glucose

Suppl. movie 8 - villus epithelium activation by acetate

Suppl. movie 9 - villus epithelium activation by L-phe

Suppl. movie 11 - max projection confocal SMP-MP recording during villus depolarisation

## Acknowledgements

Grant support: METH/014/05, KU LEUVEN. Images were recorded on microscopy equipment in LENS/Cell and tissue Imaging Cluster (CIC) supported by Hercules foundation AKUL/11/37, AKUL/09/50 and FWO G.0929.15.

## Captions Supplementary Movies

**Supplementary Movie 1.** Recording of submucosal neurons responding to the mucosal perfusion of glucose (300 mM) in a full thickness jejunal preparation from a Wnt1-cre;R26R-GCaMP3 mouse. Glucose was applied for 30 s.

**Supplementary Movie 2.** Recording of myenteric neurons responding to the mucosal perfusion of glucose (300 mM) in a full thickness jejunal preparation from a Wnt1-cre;R26R-GCaMP3 mouse. Glucose was applied for 30 s.

**Supplementary Movie 3.** Recording of submucosal neurons responding to the mucosal perfusion of acetate (100 mM) in a full thickness jejunal preparation from a Wnt1-cre;R26R-GCaMP3 mouse. Acetate was applied for 30 s. Some responses in underlying myenteric neurons that are out of focus and located to the right of the submucosal ganglion appear to precede that of the submucosal neurons.

**Supplementary Movie 4.** Recording of myenteric neurons responding to the mucosal perfusion of acetate (100 mM) in a full thickness jejunal preparation from a Wnt1-cre;R26R-GCaMP3 mouse. Acetate was applied for 30 s. The response in the initial responding myenteric neuron appears to precede that of the submucosal ganglion that is out of focus and above the myenteric ganglion.

**Supplementary Movie 5.** Recording of submucosal neurons responding to the mucosal perfusion of L-phenylalanine (L-phe; 100 mM) in a full thickness jejunal preparation from a Wnt1-cre;R26R-GCaMP3 mouse. L-phe was applied for 30 s.

**Supplementary Movie 6.** Recording of myenteric neurons responding to the mucosal perfusion of L-phenylalanine (L-phe; 100 mM) in a full thickness jejunal preparation from a Wnt1-cre;R26R-GCaMP3 mouse. L-phe was applied for 30 s.

**Supplementary Movie 7.** Epithelial responses to the perfusion of glucose (300 mM) onto the mucosal surface of a full thickness jejunal preparation from a Villin-cre;R26R-GCaMP3 mouse. The tips of villi were imaged from a lens positioned above the preparation. Glucose was applied for 1 min.

**Supplementary Movie 8.** Epithelial responses to the perfusion of acetate (100 mM) onto the mucosal surface of a full thickness jejunal preparation from a Villin-cre;R26R-GCaMP3 mouse. The tips of villi were imaged from a lens positioned above the preparation. Acetate was applied for 1 min.

**Supplementary Movie 9.** Epithelial responses to the perfusion of L-phenylalanine (L-phe; 100 mM) onto the mucosal surface of a full thickness jejunal preparation from a Villin-cre;R26R-GCaMP3 mouse. The tips of villi were imaged from a lens positioned above the preparation. L-phe was applied for 1 min.

**Supplementary Movie 10.** Example recording of myenteric neurons responding to the injection of ATP (200 μM; 2 s duration) into a villus in a full thickness jejunal preparation from a Wnt1-cre;R26R-GCaMP3 mouse.

**Supplementary Movie 11.** A spinning disk confocal microscope recording of submucosal (magenta) and myenteric (green) neurons responding to mucosal depolarization (high K^+^ (75 mM) Krebs solution applied for 10 s) in a full thickness jejunal preparation from a Wnt1-cre;R26R-GCaMP3 mouse. The top panel displays a maximum projection of the Z stack (25 μm) in the XY plane and the bottom panel depicts a maximum projection in the XZ plane. Note that the response in the myenteric plexus precedes that in the submucosal plexus.

## References

Akiba Y, Inoue T, Kaji I, Higashiyama M, Narimatsu K, Iwamoto K-i, Watanabe M, Guth PH, Engel E, Kuwahara A, Kaunitz JD (2015) Short-chain fatty acid sensing in rat duodenum. The Journal of Physiology 593:585–599.

Alcaino C, Knutson KR, Treichel AJ, Yildiz G, Strege PR, Linden DR, Li JH, Leiter AB, Szurszewski JH, Farrugia G, Beyder A (2018) A population of gut epithelial enterochromaffin cells is mechanosensitive and requires Piezo2 to convert force into serotonin release. Proceedings of the National Academy of Sciences 115:E7632.

Amato A, Cinci L, Rotondo A, Serio R, Faussone-Pellegrini MS, Vannucchi MG, Mule F (2010) Peripheral motor action of glucagon-like peptide-1 through enteric neuronal receptors. Neurogastroenterology and Motility 22:664–e203.

Bellono NW, Bayrer JR, Leitch DB, Castro J, Zhang C, O’Donnell TA, Brierley SM, Ingraham HA, Julius D (2017) Enterochromaffin Cells Are Gut Chemosensors that Couple to Sensory Neural Pathways. Cell 170:185–198.e116.

Bertrand PP (2009) The Cornucopia of Intestinal Chemosensory Transduction. Frontiers in Neuroscience 3:48.

Bertrand PP, Bornstein JC (2002) ATP as a putative sensory mediator: activation of intrinsic sensory neurons of the myenteric plexus via P2X receptors. Neuroscience 22:4767–4775.

Bertrand PP, Kunze WA, Bornstein JC, Furness JB (1998) Electrical mapping of the projections of intrinsic primary afferent neurones to the mucosa of the guinea-pig small intestine. Neurogastroenterology and Motility 10:533–541.

Bertrand PP, Kunze WA, Bornstein JC, Furness JB, Smith ML (1997) Analysis of the responses of myenteric neurons in the small intestine to chemical stimulation of the mucosa. American Journal of Physiology 273:G422–435.

Bertrand PP, Kunze WA, Furness JB, Bornstein JC (2000) The terminals of myenteric intrinsic primary afferent neurons of the guinea-pig ileum are excited by 5-hydroxytryptamine acting at 5-hydroxytryptamine-3 receptors. Neuroscience 101:459–469.

Boesmans W, Hao MM, Vanden Berghe P (2017) Optogenetic and chemogenetic techniques for neurogastroenterology. Nature Reviews Gastroenterology & Hepatology 15:21.

Chalazonitis A, Rao M (2018) Enteric nervous system manifestations of neurodegenerative disease. Brain Research 1693:207–213.

Cooke HJ, Wunderlich J, Christofi FL (2003) “The force be with you”: ATP in gut mechanosensory transduction. News in Physiological Sciences 18:43–49.

Cryan JF, O’Riordan KJ, Cowan CSM, Sandhu KV, Bastiaanssen TFS, Boehme M, Codagnone MG, Cussotto S, Fulling C, Golubeva AV, Guzzetta KE, Jaggar M, Long-Smith CM, Lyte JM, Martin JA, Molinero-Perez A, Moloney G, Morelli E, Morillas E, O’Connor R, Cruz-Pereira JS, Peterson VL, Rea K, Ritz NL, Sherwin E, Spichak S, Teichman EM, van de Wouw M, Ventura-Silva AP, Wallace-Fitzsimons SE, Hyland N, Clarke G, Dinan TG (2019) The Microbiota-Gut-Brain Axis. Physiol Rev 99:1877–2013.

Danielian PS, Muccino D, Rowitch DH, Michael SK, McMahon AP (1998) Modification of gene activity in mouse embryos in utero by a tamoxifen-inducible form of Cre recombinase. Current Biology 8:1323–1326.

Depoortere I (2014) Taste receptors of the gut: emerging roles in health and disease. Gut 63:179–190.

Diwakarla S, Fothergill LJ, Fakhry J, Callaghan B, Furness JB (2017) Heterogeneity of enterochromaffin cells within the gastrointestinal tract. Histochemistry and Cell Biology 29.

Drokhlyansky E, Smillie CS, Van Wittenberghe N, Ericsson M, Griffin GK, Eraslan G, Dionne D, Cuoco MS, Goder-Reiser MN, Sharova T, Kuksenko O, Aguirre AJ, Boland GM, Graham D, Rozenblatt-Rosen O, Xavier RJ, Regev A (2020) The Human and Mouse Enteric Nervous System at Single-Cell Resolution. Cell 182:1606–1622.e1623.

el Marjou F, Janssen KP, Chang BH, Li M, Hindie V, Chan L, Louvard D, Chambon P, Metzger D, Robine S (2004) Tissue-specific and inducible Cre-mediated recombination in the gut epithelium. Genesis (New York, NY: 2000) 39:186–193.

Ellis M, Chambers JD, Gwynne RM, Bornstein JC (2013) Serotonin and cholecystokinin mediate nutrient-induced segmentation in guinea pig small intestine. American Journal of Physiology Gastrointestinal and Liver Physiology 304:G749–761.

Foong JPP, Tough IR, Cox HM, Bornstein JC (2014) Properties of cholinergic and non-cholinergic submucosal neurons along the mouse colon. Journal of Physiology 592:777–793.

Fothergill LJ, Furness JB (2018) Diversity of enteroendocrine cells investigated at cellular and subcellular levels: the need for a new classification scheme. 150:693–702.

Fung C, Ellis M, Bornstein JC (2010) Luminal Cholera Toxin Alters Motility in Isolated Guinea-Pig Jejunum via a Pathway Independent of 5-HT(3) Receptors. Frontiers in Neuroscience 4:162.

Fung C, Vanden Berghe P (2020) Functional circuits and signal processing in the enteric nervous system. Cell Mol Life Sci. 2020 Nov;77(22):4505–4522.

Furness JB (2012) The enteric nervous system and neurogastroenterology. Nature Reviews Gastroenterology & Hepatology 9:286–294.

Furness JB, Jones C, Nurgali K, Clerc N (2004) Intrinsic primary afferent neurons and nerve circuits within the intestine. Progress in Neurobiology 72:143–164.

Gribble FM, Reimann F (2016) Enteroendocrine Cells: Chemosensors in the Intestinal Epithelium. Annual Review of Physiology 78:277–299.

Gribble FM, Reimann F (2019) Function and mechanisms of enteroendocrine cells and gut hormones in metabolism. Nature Reviews Endocrinology 15:226–237.

Gwynne RM, Bornstein JC (2007a) Local inhibitory reflexes excited by mucosal application of nutrient amino acids in guinea pig jejunum. American Journal of Physiology Gastrointestinal and Liver Physiology 292:G1660–1670.

Gwynne RM, Bornstein JC (2007b) Mechanisms underlying nutrient-induced segmentation in isolated guinea pig small intestine. American Journal of Physiology - Gastrointestinal and Liver Physiology 292:G1162–G1172.

Gwynne RM, Thomas EA, Goh SM, Sjovall H, Bornstein JC (2004) Segmentation induced by intraluminal fatty acid in isolated guinea-pig duodenum and jejunum. Journal of Physiology 556:557–569.

Haber AL, Biton M, Rogel N, Herbst RH, Shekhar K, Smillie C, Burgin G, Delorey TM, Howitt MR, Katz Y, Tirosh I, Beyaz S, Dionne D, Zhang M, Raychowdhury R, Garrett WS, Rozenblatt-Rosen O, Shi HN, Yilmaz O, Xavier RJ, Regev A (2017) A single-cell survey of the small intestinal epithelium. Nature 551:333–339.

Hao MM, Fung C, Boesmans W, Lowette K, Tack J, Vanden Berghe P (2020) Development of the intrinsic innervation of the small bowel mucosa and villi. American Journal of Physiology-Gastrointestinal and Liver Physiology 318:G53–G65.

Kaelberer MM, Buchanan KL, Klein ME, Barth BB, Montoya MM, Shen X, Bohórquez DV (2018) A gut-brain neural circuit for nutrient sensory transduction. Science 361:eaat5236.

Li Z, Fung C, Vanden Berghe P (2020) Electric Activity and Neuronal Components in the Gut Wall., 2020. In: Kuipers, EJ (ed.) Encyclopedia of Gastroenterology, 2nd edition, pp. 133–145. Oxford: Academic Press.

Li Z, Hao MM, Van den Haute C, Baekelandt V, Boesmans W, Vanden Berghe P (2019) Regional complexity in enteric neuron wiring reflects diversity of motility patterns in the mouse large intestine. eLife 8:e42914.

Liddle RA (2018) Interactions of Gut Endocrine Cells with Epithelium and Neurons. Comprehensive Physiology 8:1019–1030.

Mao Y, Wang B, Kunze W (2006) Characterization of Myenteric Sensory Neurons in the Mouse Small Intestine. Journal of Neurophysiology 96:998–1010.

Martin AM, Lumsden AL, Young RL, Jessup CF, Spencer NJ, Keating DJ (2017a) The nutrient-sensing repertoires of mouse enterochromaffin cells differ between duodenum and colon. 29.

Martin AM, Lumsden AL, Young RL, Jessup CF, Spencer NJ, Keating DJ (2017b) Regional differences in nutrient-induced secretion of gut serotonin. Physiological Reports 5:e13199.

Melo CGS, Nicolai EN, Alcaino C, Cassmann TJ, Whiteman ST, Wright AM, Miller KE, Gibbons SJ (2020) Identification of intrinsic primary afferent neurons in mouse jejunum. e13989.

Mongardi Fantaguzzi C, Thacker M, Chiocchetti R, Furness JB (2009) Identification of neuron types in the submucosal ganglia of the mouse ileum. Cell and Tissue Research 336:179–189.

Moore BA, Vanner S (2000) Properties of synaptic inputs from myenteric neurons innervating submucosal S neurons in guinea pig ileum. American Journal of Physiology Gastrointestinal and Liver Physiology 278:G273–280.

Obata Y, Pachnis V (2016) The Effect of Microbiota and the Immune System on the Development and Organization of the Enteric Nervous System. Gastroenterology 151:836–844.

Qu ZD, Thacker M, Castelucci P, Bagyánszki M, Epstein ML, Furness JB (2008) Immunohistochemical analysis of neuron types in the mouse small intestine. Cell and Tissue Research 334:147–161.

Reitelseder S, Tranberg B, Agergaard J, Dideriksen K, Højfeldt G, Merry ME, Storm AC, Poulsen KR, Hansen ET, van Hall G, Lund P, Holm L (2020) Phenylalanine stable isotope tracer labeling of cow milk and meat and human experimental applications to study dietary protein-derived amino acid availability. Clinical Nutrition.

Richards P, Parker HE, Adriaenssens AE, Hodgson JM, Cork SC, Trapp S, Gribble FM, Reimann F (2014) Identification and characterisation of glucagon-like peptide-1 receptor expressing cells using a new transgenic mouse model. Diabetes 63:1224–1233.

Rogers GJ, Tolhurst G, Ramzan A, Habib AM, Parker HE, Gribble FM, Reimann F (2011) Electrical activity-triggered glucagon-like peptide-1 secretion from primary murine L-cells. The Journal of physiology 589:1081–1093.

Schemann M, Ehrlein HJ (1986) Postprandial patterns of canine jejunal motility and transit of luminal content. Gastroenterology 90:991–1000.

Schneider S, Wright CM, Heuckeroth RO (2019) Unexpected Roles for the Second Brain: Enteric Nervous System as Master Regulator of Bowel Function. Annual Review of Physiology 81:235–259.

Spencer NJ, Hu H (2020) Enteric nervous system: sensory transduction, neural circuits and gastrointestinal motility. Nature Reviews Gastroenterology & Hepatology.

Su CY, Menuz K, Carlson JR (2009) Olfactory perception: receptors, cells, and circuits. Cell 139:45–59.

Vanden Berghe P, Kenyon JL, Smith TK (2002) Mitochondrial Ca2+ uptake regulates the excitability of myenteric neurons. Journal of Neuroscience 22:6962–6971.

Ventura EE, Davis JN, Goran MI (2011) Sugar content of popular sweetened beverages based on objective laboratory analysis: focus on fructose content. Obesity (Silver Spring) 19:868–874.

Wang Y, DelRosso NV, Vaidyanathan TV, Cahill MK, Reitman ME, Pittolo S, Mi X, Yu G, Poskanzer KE (2019) Accurate quantification of astrocyte and neurotransmitter fluorescence dynamics for single-cell and population-level physiology. Nature neuroscience 22:1936–1944.

Wong V, Blennerhassett M, Vanner S (2008) Electrophysiological and morphological properties of submucosal neurons in the mouse distal colon. Neurogastroenterology and Motility 20:725–734.

Zariwala HA, Borghuis BG, Hoogland TM, Madisen L, Tian L, De Zeeuw CI, Zeng H, Looger LL, Svoboda K, Chen TW (2012) A Cre-dependent GCaMP3 reporter mouse for neuronal imaging in vivo. Journal of Neuroscience 32:3131–3141.

Zeisel A, Hochgerner H, Lonnerberg P, Johnsson A, Memic F, van der Zwan J, Haring M, Braun E, Borm LE, La Manno G, Codeluppi S, Furlan A, Lee K, Skene N, Harris KD, Hjerling-Leffler J, Arenas E, Ernfors P, Marklund U, Linnarsson S (2018) Molecular Architecture of the Mouse Nervous System. Cell 174:999–1014.e1022.

Zhou X, Galligan JJ (1999) Synaptic activation and properties of 5-hydroxytryptamine(3) receptors in myenteric neurons of guinea pig intestine. The Journal of Pharmacology and Experimental Therapeutics 290:803–810.

